# Salinity is associated with habitat use and mitochondrial lineage diversity in halophilic *Ephydra* flies (Diptera: Ephydridae)

**DOI:** 10.64898/2026.09.03.749233

**Authors:** Kirsten I. Verster, David Herbst, Ekaterina Yakovleva, Sarah Borja, Emily Celallos Fuentes, Vianda Nguyen, Andrew Wang, Ashley Wong, Alina Zhang, Carly Biedul, Bonnie Baxter

## Abstract

Brine flies of the genus *Ephydra* (Diptera: Ephydridae) are ubiquitous extremophile animals whose larval forms are able to thrive in high-stress environments, including hypersaline and high carbonate waters. Despite their historical and ecological importance, little is known about their evolutionary history. Here, we focus on an understudied population of *Ephydra* living in former industrial salt evaporation ponds of the San Francisco Bay Area (SFBA), a region which has undergone significant environmental changes in the past century. We observed and collected *Ephydra* specimens and took salinity measurements of waters they inhabited during a 6-month period in 2024. We found that *E. millbrae* and *E. gracilis* demonstrate strong salinity preferences. We also analyzed DNA from *Ephydra* spp. specimens from distant sites and a timespan of ∼40 years, generating the most comprehensive gene genealogy of the group to date based on a single mitochondrial locus. Our analysis of phylogenetic signals suggests that salinity envelopes are influenced by evolutionary relationships in *Ephydra*. This study provides new information about the natural history of extremophile flies in the SFBA and fills in current gaps in our knowledge about the evolutionary history of the genus more broadly.

## Introduction

Hypersaline water bodies are osmotically challenging environments, and yet many organisms live and thrive in them. At low salinities, maintaining homeostasis necessitates membrane proteins that use electrochemical gradients to expel excess ions in the cell (Dubyak, 2004). However, hypersaline environments represent additional osmotic stress that requires strategies such as removing intracellular sodium, which can accumulate and deactivate enzymes by interfering with cationic sites (Dennis & Shimmin, 1997).

While there is a high diversity of halophilic microbes (Oren, 2024), there are very few invertebrates that can thrive in these environments. For example, the open waters of Great Salt Lake (GSL) in Utah, USA are known to contain only three invertebrate taxa: brine shrimp (*Artemia franciscana*), brine flies (*Ephydra* spp.), and nematodes (Jung et al., 2024). Yet, that lack of invertebrate diversity is accompanied by great abundance. For example, surface aggregations of brine shrimp can be unambiguously identified from satellite imagery (Qi et al., 2021). The abundance of halophilic invertebrates may be due to the absence of competition or predation (Herbst, 2001).

Halophiles have evolved unique adaptations to survive these high-stress environments. For example, halophilic species (such as fungi, archaea and plants) have evolved changes in membrane composition to survive in hypersaline environments (Dennis & Shimmin, 1997; Turk et al., 2004), or specialized vacuoles that sequester Na+. The genetic changes underlying these adaptations are largely unknown, with some notable exceptions. For example, a recent study showed that the brine shrimp has an isoform of the otherwise highly conserved Na+/K+--ATPase, which, in the presence of high salinities, demonstrates a reduced stoichiometry - flipping the script on the fundamental rules of cellular energetics (Artigas et al., 2023). Other extremophile invertebrates may likewise have evolved unexpected, paradigm-shifting tools to survive. One such group includes the genus *Ephydra* (Diptera: Ephydridae), a genus of flies commonly known as brine flies (Herbst & Bradley, 1989).

*Ephydra* flies have been found breeding by the millions in salt-water ponds around the world and serve as an important food sources for local wildlife (Wurtsbaugh, 2009). They are critical food sources for Wilson’s phalaropes, in places like Mono Lake, CA and the GSL (Frank & Conover, 2019). They have also historically been used as food sources for Indigenous peoples such as the Paiute (Great Basin) or Kutzadika’a (from Mono Lake) (Wirth, 1971; Schrader et al., 2016). They are adapted to the most extreme aquatic chemical environments, such as hypersaline lakes (e.g. GSL), saline alkaline lakes (e.g. Mono Lake, CA), and even acidic thermal springs (e.g. Yellowstone, WY) (Wirth, 1975). Because *Ephydra* occupy diverse aquatic niches (**Fig. 1**) (Herbst, 1999), they may represent an adaptive radiation, which, in turn, lends itself to a compelling comparative framework that could address how species have evolved to colonize diverse, chemically severe environments. Despite their abundance, extreme lifestyle, and historical significance, the biology and evolutionary history of *Ephydra* brine flies remains poorly understood, especially compared to other dipteran taxa such as *Drosophila*.

**Fig. 1.**
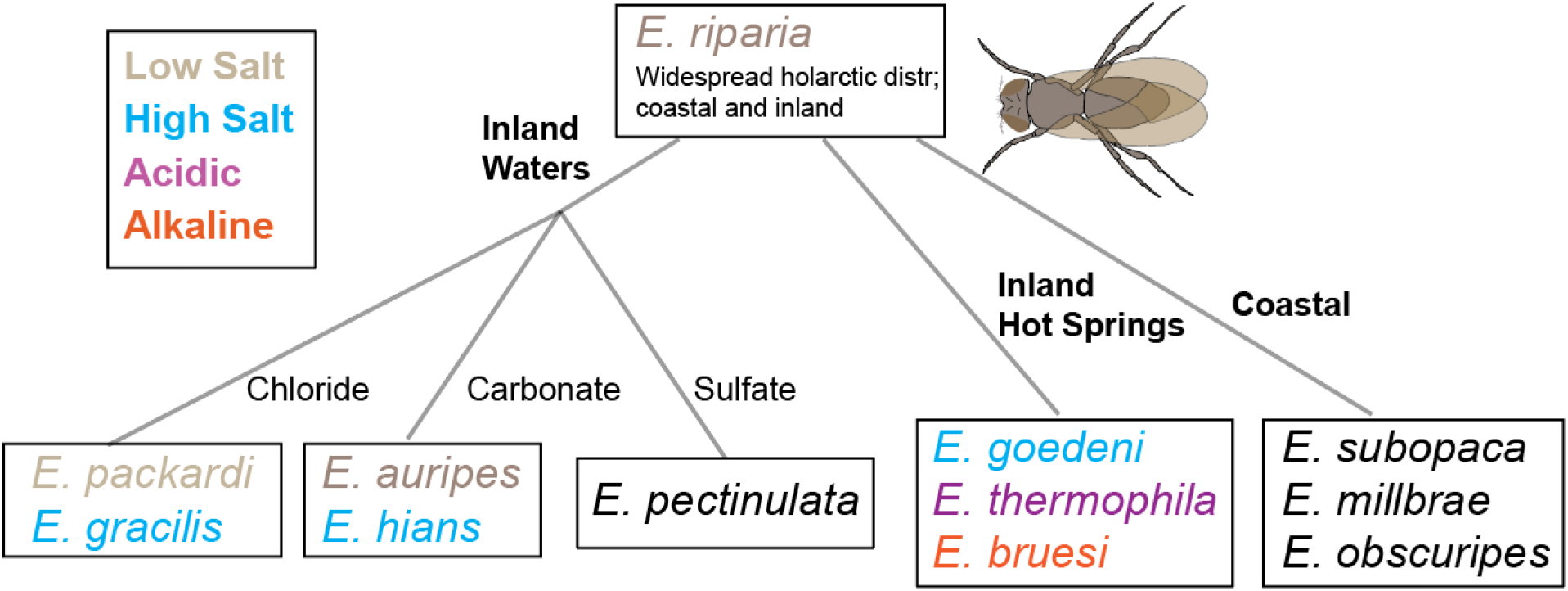
The diverse water habitat chemistries of some Nearctic *Ephydra* species. Image adapted from (Herbst, 2001).

Most studies about *Ephydra* focus on *E. gracilis* from GSL, UT (Herbst, 1999, 2023) and the “alkali fly” *E. hians* Say from Mono Lake, CA (Herbst, 1988, 1990, 1999). Other species and populations of *Ephydra* have not received as much recognition, but are locally known for their ubiquity and critical role in feeding avian communities, such as those from the South Bay Salt Ponds (SBSP) in the San Francisco Bay Area (SFBA) of California, USA (Takekawa et al., 2011).

The SBSP in California were originally natural salt marsh ecosystems utilized by the Indigenous Ohlone (Johnson, 1996). They were converted into industrial salt evaporation ponds during the Gold Rush, when salt was critical for food preservation (Brown, 1960). After European colonization of the area, the salt ponds were run by small, family-owned companies (Brown, 1960). Following the California Legislature Revising Act (March 28, 1868), they were then subsequently bought up by Oliver Salt Company, Leslie Salt Company, and eventually Cargill (Palaima, 2012). In 2003, Cargill sold the majority of its holdings to various government and state entities that later became the South Bay Salt Pond Restoration Project (SBSPRP), an initiative launched with the aim of restoring the salt ponds to tidal wetlands. As of 2019, approximately 1,300 ha of managed ponds have been breached to tidal action to begin tidal marsh restoration (US Geological Survey et al., 2021), which is expected to shift salinity over time. Currently, the majority of the former salt ponds are no longer in operation. Many species in the SBSP are adapted to saline and hypersaline environments, though few studies have been conducted exploring their biology and ecology (Carpelan, 1957). Consequently, there are large knowledge gaps when planning enhancement or restoration projects.

SBSP brine flies are important food for several other species, such as coleopteran *Tanarthus occidentalis* and *Cicindela senilis senilis*, dictynid and salticid spiders, and numerous bird species including snowy plovers, California gulls, black necked stilts, avocets, and more (Maffei, 2000). There are two known brine fly species in the SBSP, *E. millbrae* Jones and *E. gracilis* Packard (Maffei, 2000).

*E. millbrae* were so named given that they were originally found near Millbrae, CA in 1906 (Jones, 1906). They are a blue-green species, initially described as “very abundant along the southwest shore [of the SFBA]… the floating puparia and adults often cover the entire surface of the small salt-water ponds” (Jones, 1906). *E. gracilis* is one of the abundant fly species found from the GSL, though they have also been identified in the southwestern United States and even Hawaii. They are known to occur in tremendous numbers in high chloride, coastal and inland salt waters, including the SBSP. Of the two, *E. millbrae* is far less studied.

For this study, we closed critical gaps in our understanding of the biology and natural history of *Ephydra* spp. from the SFBA. We collected abiotic and biotic data where both *E. gracilis* and *E. millbrae* were collected. Furthermore, we used *cytochrome oxidase I* (*COI*) genotyping and phylogenetic analysis to further our understanding of the evolution of these flies in the SFBA and other localities.

## Materials and Methods

### Observations of Salinity and Habitat Preferences

*Ephydra* observations and collections occurred across the SBSP (**Fig. 2a)** and other SFBA sites from March 2024 to August 2024. The presence of brine shrimp, plants, and other physical structures were noted. We gently swept various sites and collected larvae and pupae with nets, or flies with aspirators. We collected information about the salinity and pH of the water bodies from which insects were collected with a lightweight, portable large-range digital salinity and pH meter (ORAPXI, ASIN: B08GC7JCV3) that enabled us to take measurements while at the collection site (all reported salinities in the manuscript, besides PQ772809.1, were measured with this meter). Prior to fieldwork, the salinity meter was calibrated against the Sper salinity refractometer (Carolina #652388) to confirm values were accurate (within 2%) at the expected salinity ranges (20ppt-250 ppt), while pH was calibrated against Universal pH Indicator Strips (Carolina #893830). Measurements were taken twice within 2 minutes of each other at the sampling site; if not identical, the mean of the two values was written.

**Fig. 2.**
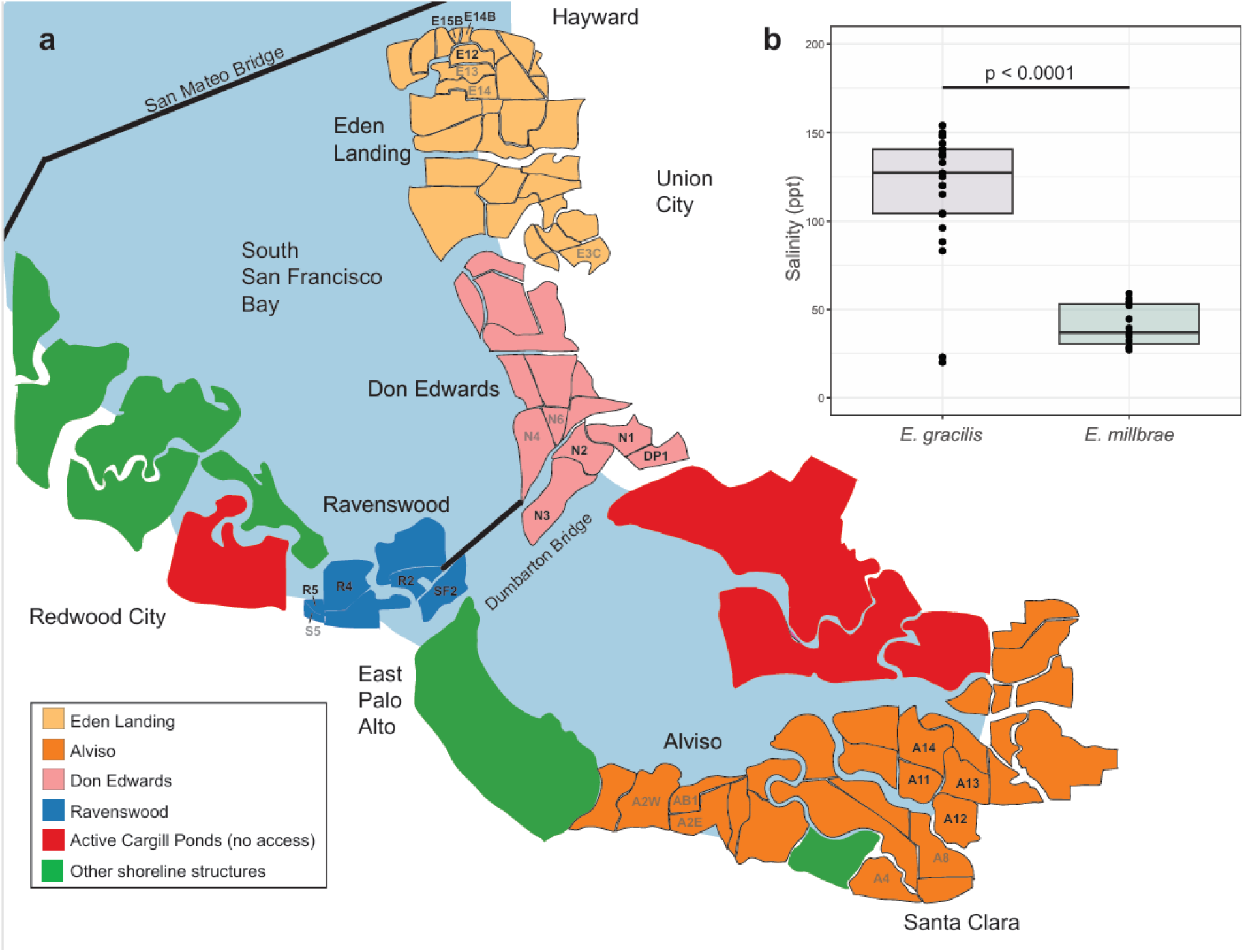
**(a)** Schematic of the SBSP pond system annotated with entities governing the different pond complexes. Grey text labels indicates the pond was visited during the study period but no flies were found, while black text labels indicates that flies of either species were found in the pond. Salinity and brine fly data for each observed pond can be found in **Supplementary Table 1. (b)** *E. gracilis* and *E. millbrae* show marked preferences for non-overlapping salinity envelopes, with *E. gracilis* and *E. millbrae* showing preferences for salinities at ∼80-150 ppt and ∼30-60 ppt, respectively.

Adult representative specimens were initially identified to species-level via genital dissection (Wirth, 1971), after which striking differences in size and coloration were used to discriminate between the two SFBA species. Larval and pupal stages were confirmed to match with the adult flies via *COI* genotyping, after which striking differences in morphology were used to discriminate between species. All species identifications were confirmed by the lead investigator of this study (KIV).

We visited and took measurements from 26 named SFBA former salt ponds over 13 separate days, approximately once a week, from March 2024 to the end of August 2024. The unique named ponds are from the definitive SBSPR pond ID system (finalized in 2011). For our observations of *Ephydra* spp. preferences, we included ponds for which we identified *at least* five living *Ephydra* of the given species at any life stage, either in or immediately adjacent to the water body. Brine shrimp were marked as present if more than five living hatched shrimp were found.

Flies were collected under the following sampling permission for the SBSP: Santa Clara County Parks 1643956, East Bay Regional Parks District 23-1194, and Don Edwards San Francisco Bay National Wildlife Refuge 2023-14. Additional piecemeal sampling was conducted under permits #25-02 (Mono Lake Tufa State Natural Reserve) and Right of Entry Permit No. 410-00807 (GSL), with written permission for other sites.

### Phylogenetic Analysis

#### DNA Extraction

DNA was extracted from single individual flies using the QIAGEN DNeasy Blood & Tissue Kit animal tissue protocol (modified to include a shaking overnight incubation with microfuge vials that were laid horizontally to promote mixing). Specimens collected from the SBSP were frozen at -80°C the day of collection before processing. Specimens from all other sites, including samples preserved in ethanol that were over 40 years old, were preserved at room temperature in 70% or 100% EtOH and then left out to dry on a Kimwipe for ∼10 m prior to DNA extraction. While DNA was extracted from pinned *Ephydra* specimens from Cal Academy of Sciences (San Francisco, CA) and Essig Museum (Berkeley, CA), we were unable to extract amplifiable DNA from these collections.

#### PCR, Gel Visualization, and Sequencing

PCR reaction mixes were done following GoTaq ® Green Master Mix (Promega) protocols using 25-200 ng of gDNA from single flies in reaction volumes of 15 µl. The mitochondrial gene *COI* was amplified using primers BF2 (5’-GCHCCHGAYATRGCHTTYCC-3’) and BR2 (5’-TCDGGRTGNCCRAARAAYCA-3’), which were chosen since they outperform traditionally used Folmer barcoding primers in aquatic insects (Elbrecht & Leese, 2017). Thermal cycler settings were: 5 m at 95°C and 35 cycles of: 95°C for 30 s, 50°C for 30 s, and 68°C for 30 s, followed by a final 5 m extension at 68°C. 1% agarose 1X TBE gels were prepared in 1X TBE buffer with 1 µL SYBR™ Safe staining gel per 10 mL of gel solution. 4 µL of PCR product was mixed with 1 µL ThermoScientific 6X Loading Dye and 100bp DNA ladder (NEB) was included as a molecular marker. PCR product was run on gels using the Owl™ EasyCast™ B1 Mini Gel Electrophoresis System rigs for 15-25 minutes at 120V and visualized in a Gel Doc XR+ (Bio-Rad). PCR amplicons and primers were submitted to ElimBio for PCR cleanup and Sanger sequencing. Forward and reverse sequences were trimmed, merged, and analyzed with Geneious Prime v.2024.0.5. Heterozygous nucleotides were manually annotated. Sequences are deposited in GenBank under accession IDs PQ772805-PQ772817, PV790562-PV790574, and PZ634866.

#### Phylogenetic Analyses

##### Alignment

We collected *COI* sequences from collected specimens, representative sequences from NCBI GenBank, and three species that served as outgroups (the lepidopteran species *Maniola jurtina* and two dipteran outgroups, *Hydrellia griseola* and *Drosophila punjabensis*). To avoid redundancy in the phylogeny, we included no more than one specimen per phylogenetic clade per collection location per year (separate ponds in the SBSP system were considered separate collection locations).

Initial iterations of this analysis were confounded by artifacts which, in retrospect, were likely nuclear mitochondrial DNA segments (NUMTs). In analyzing *Ephydra* genomes (GenBank assembly IDs GCA_001014675.1 and GCA_001015075.1), we found several BLAST hits with high percent identity to ephydrid *COI*, but which did not encode full-length ORFs and were located on long nuclear scaffolds. This suggests the presence of NUMTs (Xue et al., 2023), which can confound phylogenetic analyses (Song et al., 2008). In order to exclude potential NUMTs, ORFs were predicted using the Geneious “Find ORFs” function with a minimum size of 400 bp, including interior ORFs and continuing outside the sequence using the Invertebrate Mitochondrial genetic code (trans_table 5). Sequences which did not meet these criteria in the expected reading frame were discarded. Sequences were aligned with MAFFT v. 7.490 (Katoh & Standley, 2013) in Geneious Prime v.2024.0.5. The final alignment is shown in **Online Resource 1**.

##### Phylogenetic Construction

The maximum-likelihood gene phylogeny was generated from 39 sequences with 682 nucleotide sites from IQTree v. 3.1.2 (Wong et al., 2026) (the newick file can be found in **Online Resource 2**). Branch support was assessed using 1,000 ultrafast bootstrap (UFBoot) replicates. The phylogeny was rooted with the butterfly *Maniola jurtina*. We collapsed nodes with < 90% UFBoot support for visual clarity. The tree was annotated with ggtree v3.12.0 (Yu et al., 2017) in RStudio 2024.4.1.748. Species for which salinity data were collected at the larval or pupal stage (*n*=17) were annotated with the ggtree function “gheatmap”. Unlike our SBSP natural history analysis, only larvae and pupae were included in this analysis since their presence in the water body suggests some degree of physiological tolerance, whereas an adult fly could stumble near a shoreline by chance.

##### Calculation of Phylogenetic Signal

We used the “phylosig” function in the R package phytools (Revell, 2024) to calculate Blomberg’s K (Blomberg et al., 2003) and Pagel’s λ (Pagel, 1999) for salinity. The tree was pruned to 17 tips (the number for which larval and pupal salinity had been measured) for the analysis.

## Results

### Natural History and Habitat Preference of San Francisco Bay Area Brine Flies

#### Brine fly salinity preference

We began exploratory fieldwork in March 2024, during which we identified the best ways to access the various named salt ponds as well as safe sampling sites, measured salinity and pH of these ponds, and determined which places contained invertebrate fauna and would thus be good candidate sites for recurring collections. We frequently observed heterogeneity in salinity caused by levees, ephemeral ponds, or other structures, which created distinct water profiles within the same named pond. Measured salinities of all sites ranged from 1-154 ppt and pHs ranged from 6.56 to 9.6. pH and salinity were negatively correlated (adjusted R^**2**^ = 0.7376, p-value = 3.541e-15) at the sites measured (*n=* 48 observations; data for this analysis found in **Online Resource 3**). After researchers were trained to ID species in the field, we began taking observations about salinity preferences for both *E. millbrae* and *E. gracilis*. The two species showed marked salinity preferences (Welch two sample t-test, df=28.285, p-value 3.228e-10) (**Fig 2b;** underlying data can be found in **Online Resource 4**).

#### E. gracilis *habitat preferences*

*E. gracilis* larvae and pupae (**Fig 3a**) generally occupied salinity envelopes from ∼80-150 ppt **(Fig 2b)**, though some were found in ponds with salinities as low as 20 ppt (these were within walking distance of ponds of higher salinity, so it is possible that they travelled from nearby). Where *E. gracilis* were typically found, vegetation along the ponds were scarce (e.g. **Fig 3b**). Overlapping cohorts of larvae, pupae, and adult flies were found during the time sampled. Larvae were found free-floating in salt ponds or crawling on sediment or substrate. Pupae were found attached upright to underwater substrates such as rocks (**Fig 3c**), littered along the shoreline (e.g. **Fig 3d**), or in some cases appeared to be attached to hard clumps of mud at the bottom of still salt ponds. Adults tended to prefer muddy areas along salt ponds and would often appear in large swarms (**Fig 3e**). When *E. gracilis* (of any life stage) were found alive in a given salt pond, living brine shrimp were found 61.9% of the time (**Online Resource 3**, *n*=21 observations).

**Fig. 3.**
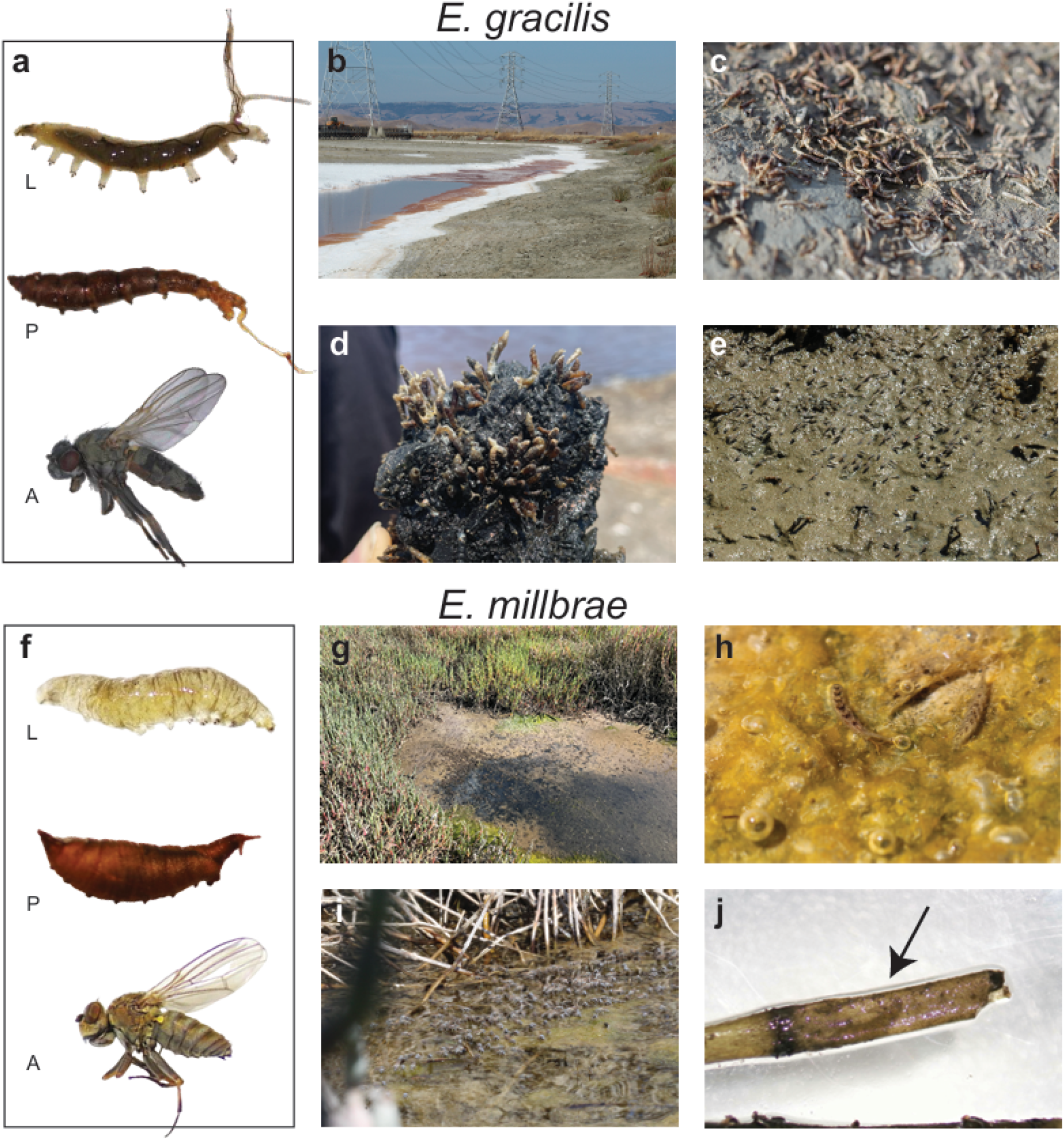
Photos highlighting the morphology across development and preferred habitats of *E. gracilis* (a-e) or *E. millbrae* (f-j) (described in the text). Acronyms in a, f: L = larvae; P = pupae; A=adult.

#### E. millbrae *habitat preferences*

*E. millbrae* (**Fig 3f**) were primarily found in marsh tidal pools ∼30-60 ppt **(Fig 2b)**. Sites in which they were found were frequently lined with vascular vegetation such as pickleweed and *Ulva* sp. (**Fig 3g**), in addition to microbial mats (**Fig 3h**). Despite finding large swarms of adult flies (**Fig 3i)**, it was initially difficult to find larvae and pupae; however, eventually both were found deeply embedded in unidentified decaying vegetation (**Fig 3j**) or enwrapped in *Ulva* sp. in ambient bay water. We never found *E. millbrae* in the same pond as brine shrimp (**Online Resource 3**, *n*=15 observations).

### Phylogenetic Analysis of Brine Flies

The maximum likelihood gene genealogy is shown in **Fig 4** (best-fit model according to BIC: GTR+F+G4; LogL = -3132.302; BIC=6813.320). The tree shows a highly-supported *E. hians* clade **[A]**, which includes samples from across the Great Basin and adjacent western North America, including Oregon (Lake Abert), Colorado (Big Soda Lake), Utah (GSL), and California (Owens Lake, Mono Lake, Deep Springs) and spanning collection dates from 1984 to the present. Another clade [**B]** includes several species from the Riparia group (e.g. *E. macellaria* Egger, *E. packardi* Wirth, and *E. riparia* Fallén (Wirth, 1971)), as well as several unidentified species from diverse localities including Utah (USA) and Canada. In our tree, *E. millbrae* [**C**] is sister to clade [**B]** (with 91% UFBoot Support, though we note that UFBoot support is optimal at >95% according to convention (Hoang et al., 2018)). We show the tree at a 90% cutoff for visual clarity and since *E. millbrae* is considered to be part of the Riparia group (Wirth, 1971). On our gene genealogy, *E. gracilis* is paraphyletic: one lineage includes flies from various populations in the SBSP, and another from the GSL (Haney et al., 2023).

**Fig. 4.**
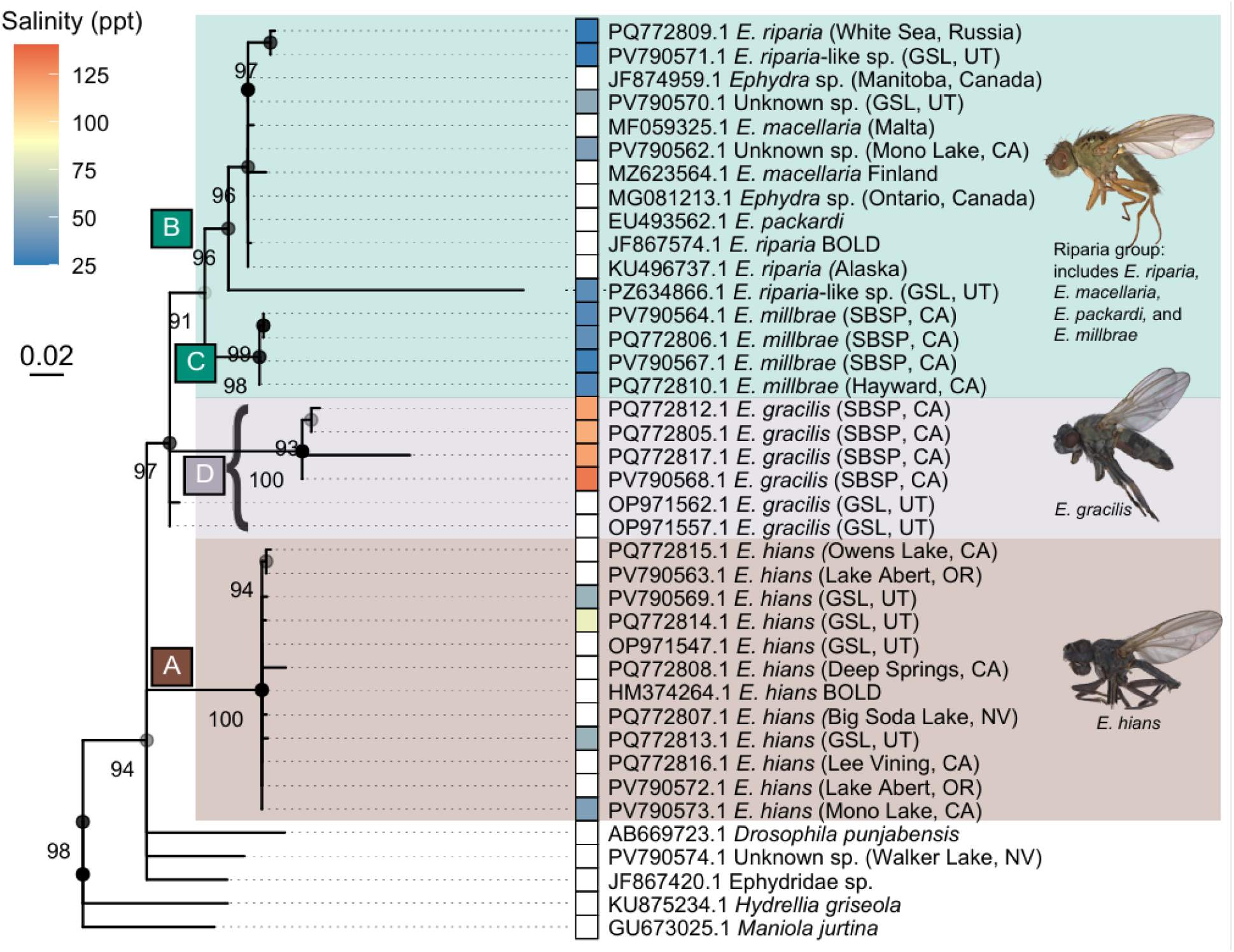
Annotated *COI* gene genealogy of *Ephydra* spp.; node values represent UFBoot (%). Acronyms: SBSP = South Bay Salt Ponds; GSL = Great Salt Lake. Annotated clades [A-D] are described in the text.

For our phylogenetic signal calculations of salinity, we measured Blomberg’s K and Pagel’s λ. Pagel’s λ was 0.925 (logL = -73.467, LR = 21.310, p-value based on LR test = 3.90e-6). Blomberg’s K was 0.024 (p-value based on 1000 randomizations = 0.002). The results, coupled with our annotations (**Fig 4**), suggests that some closely related species may be ecologically different (low K), but overall the salinity habitats are phylogenetically structured (high λ).

## Discussion

### Salinity Preferences of SBSP *Ephydra*

Our natural history observations of *E. millbrae* and *E. gracilis* in the SBSP extends our knowledge of ephydrids in this region in the present day, at a time when the SBSPs are experiencing significant abiotic and biotic changes due to anthropogenic interventions. *Ephydra* were listed as candidate sentinel species for monitoring mercury in the region, and subsequent analysis showed that they indicate mercury bioaccumulation in the salt panne food web (Kieu et al, 2009). Lack of knowledge about their basic biology can lead to a failure to utilize them effectively to better understand Bay Area ecology; for example, a previous study examining pesticide safety in nontarget insects was unable to collect data on *E. gracilis* emergence, likely due to their being placed in non-optimal salinities (Lawler et al., 2000). Marked habitat heterogeneity (e.g., salt ponds of ∼150 ppt could be within 25 feet of ponds ∼30 ppt) may have complicated previous assessments of *Ephydra* distributions. Understanding the biology of these flies, which play a vital role in salt pond ecosystems as a food source for avian species (Takekawa et al., 2006; Frank & Conover, 2023), is critical as the landscape undergoes anthropogenic interventions.

### Phylogenetic Analyses of *Ephydra*

#### Topology

In our tree, nodes with <90% bootstrap support were collapsed to polytomies, revealing two major clades. The first clade predominantly features *E. hians* from across the continental USA (A), while the second clade (B + C+ D) includes species from the Riparia group and *E. gracilis*. A dated phylogeny of Drosophilidae dates this split at approximately 13.3 mya (Dias et al., 2025).

Clade **[B + C]** includes species from the Riparia group (Wirth, 1971), which also includes *E. goedeni* and *E. subopaca*. Flies from this group are considered less salt tolerant (Wirth, 1971; Herbst, 2001). *E. millbrae* forms a subclade sister to another which includes *E. macellaria, E. riparia*, and *E. packardi*, the three of which have near-identical sequence similarity at *COI* (99.4%) (*COI* similarity data included in **Online Resource 5**). This is unexpected since, in Diptera, distinct species are expected to have at least 1-3% divergence (Renaud et al., 2012; Nzelu et al., 2015). It is possible these are different morphs of the same species; some North American taxonomists considered species in this group to be varieties, such as *E. riparia* and *E. macellaria*, which were considered to be “dark-legged” and “pale-legged” morphs. However, a subsequent review of the Riparia group (Wirth, 1971) suggested 6 separate distinct species with well-correlated distributional and habitat distinctions. Indeed, *E. macellaria* is known as a rice pest (Kholmurodov & Avazov, 2024), *E. packardi* is known to prefer low saline chloride habitats (Herbst, 1999), and *E. riparia* has a widespread holarctic distribution, occupying both coastal and inland habitats (Herbst, 2001). Instead, we speculate this is the result of mitochondrial introgression (Sloan et al., 2017), incomplete lineage sorting (Maddison & Knowles, 2006), a *Wolbachia*-driven mitochondrial sweep (Cariou et al., 2017), or a combination therein. *Wolbachia* have been identified in *E. hians* and *E. gracilis* from the GSL with low sequence divergence (Truong et al., 2017), marking the last two options as a possibility.

Riparia group flies were found both in Mono Lake (PV790562.1) and GSL (PV790571.1). These likely represent *E. packardi*, which have been identified in these locations (Wirth, 1971). One group of flies from the GSL is sister to the aforementioned lineage (PZ634866.1); these were found in borrow ditches along the lake (35 ppt) and could potentially represent *E. pectinulata*, which have been previously identified in that area, has a developmental optima near that salinity (Wirth, 1971), and has no sequence data publicly available.

Monophyly of *E. gracilis* was weakly supported (64%) and collapses into a polytomy, leaving the SBSP and GSL lineages unresolved with respect to one another. We did not find *E. gracilis* in GSL during the sampling period; those included in the phylogeny were taken from existing NCBI GenBank accessions. Though they were morphologically identical to GSL *E. gracilis* based on genital dissection, *E. gracilis* from the SFBA showed high genetic divergence from GSL flies (e.g. two representative tips from the group have ∼10% divergence at *COI*), suggesting cryptic speciation.

Flies related to *E. hians* form a separate clade **[A]**. Some samples, such as those from Lake Abert in Oregon (PV790563.1), were collected over 40 years ago. Other samples were collected from various alkaline lakes across the northwestern USA (e.g. Big Soda Lake in Nevada, Deep Springs in California).

#### Phylogenetic signal

Our calculations of phylogenetic signal revealed statistically significant results, with a low estimate for Blomberg’s K (K=0.024) and high estimate for Pagel’s λ (λ=0.925). Per (Ollier et al., 2006; Münkemüller et al., 2012), we consider these results along with the annotated phylogeny (**Fig 4**). Low phylogenetic signal, as found in our Blomberg’s K calculation, can in some contexts be attributed to high evolutionary lability (Blomberg et al., 2003; Kamilar & Cooper, 2013), which is vital to adaptive radiations. Blomberg’s K is more sensitive to similarity within closely-related species (Blomberg et al., 2003). In our phylogeny, we see that there is some variation between habitats where similar species were found (e.g. in clade (A) salinity ranges from 45 ppt-83 ppt). Our high calculation of Pagel’s λ suggests the distribution of salinity envelopes are influenced by their evolutionary relationship, as we can see in **Fig 4** (e.g. the observed range in this study for species in the *E. hians* [A], *E. gracilis* [D], and Riparia [B+C] clades are 45-83 ppt, 116-133 ppt, and 25-50 ppt, respectively).

The within-clade variance of Blomberg’s K may represent some of the spatiotemporal stochasticity of the measurement we are applying. Habitat salinity is not a fixed physiological trait like body size but is inherently a fluctuating measurement that can change over the course of a day or season in the same place. However, we can assume this salinity corresponds to some combination of the clades’s physiological optima as well as preference. While some studies have specifically analyzed the physiological impact of salinity on *Ephydra* (Herbst, 1999, 2023), these experiments have not been conducted across all *Ephydra* species and so we include individual specimens collection data as opposed to mean trait values. The relatively low number of tips included in the analysis could impact our calculations (Freckleton et al., 2002), though polytomies are expected to have negligible effects on the estimates of phylogenetic signal (Münkemüller et al., 2012). Furthermore, other important aspects of water chemistry, such as carbonate concentration, are missing from this analysis and may further explain habitat partitioning in *Ephydra*.

#### Implications for species barcoding

In initial implementations of this study, Sanger sequencing occasionally resulted in *COI* sequences that appeared to have frameshift mutations, stop codons, or encode large deletions. We later determined these were NUMT artifacts complicating our analysis and removed them from our alignment. Correspondingly, previous studies have shown NUMTs can lead to overestimation of species richness (Song et al., 2008). While beyond the scope of this study, we speculate that hypersaline environments might support NUMT formation. While desiccation-prone environments have not been shown to increase NUMT formation(Li et al., 2020), it is possible other factors in the *Ephydra* environment, such as heavy metal contamination or osmotic stress, could lead to DNA damage that promotes NUMTs accumulation.

The presence of NUMTs, coupled with the other unusual patterns we see with respect to *COI* (as described above), should provoke some measure of caution in using this gene in metabarcoding, especially as its usage continues to increase in biodiversity assessments (Salis et al., 2024).

Our phylogeny reveals unexpected patterns, and generates more questions, about the evolution of this genus. While *COI* is an important gene for species identification and has been widely used for DNA barcoding (De et al., 2014), its suitability in constructing species trees can be limited (Nelson et al., 2012; Cameron, 2014; McDonagh et al., 2016). Errors in the phylogeny would, in turn, affect our interpretation of phylogenetic signal. These limitations could explain some of the unusual patterns discussed here.

To our knowledge we have assembled the most taxonomically and geographically comprehensive *COI* genealogy of *Ephydra* assembled to date, with over 6 named species, 5 countries, and >40 years of collections. However, this is far from representing the entirety of at least 29 *Ephydra* species reported (Wirth, 1971, 1975). More comprehensive sampling, coupled with sequencing of more loci (or, ideally, a phylogenomics approach), is vital to understanding the evolutionary biology of the genus.

## Conclusion

*Ephydra* flies thrive in hypersaline aquatic habitats worldwide, and play critical ecological roles globally. Many terminal saline lakes (e.g. Great Salt Lake, Mono Lake, Owens Lake, Aral Sea, and Lake Urmia) are shrinking to consumptive uses and climate change pressures. The resulting increased salinity is often exceeding the tolerance limits of brine flies and threatening their availability as food for shorebirds (Herbst, 2023). However, brine fly biology, especially that of less studied species such as *E. millbrae*, is poorly understood, particularly in the SFBA. This study closes that gap by providing new natural history data and species distributions in the SFBA. Our *COI* genealogy, which builds on the work of (Haney et al., 2023) and barcoding sequences deposited in NCBI GenBank, is currently the most comprehensive of the genus to date, laying the groundwork for future evolutionary and genomic studies of this unique, ecologically significant fly genus.

## Supporting information

Online Resources/Supplementary Material

## Acknowledgements

We thank the many park officials and colleagues who helped this research come to light. Chris Grinter for providing access to resources at Cal Academy of Sciences. Various parks officials including Karla Toledo (East Bay Parks); Carly White and Garrett Allen (Eden Landing Ecological Reserve); Rachel Tertes (Don Edwards National Wildlife Refuge); Maddy Schwarz (San Francisco Bay Bird Observatory); Dash Dunkell (Elkhorn Slough Foundation); Marisa Weinberg (Utah Division of Forestry, Fire and State Lands); Lindsay Cline (California State Parks - Sierra District). Audrey Yau, Ao Yu, Joy Kumagai, Bohdan Kamets, Marty Freeland, Sierra Reinertson, Yvette Betancourt, Jillian de la Torre, Raoul Olivier Martin, Georgie Corkery, and Pooch Martin provided fieldwork assistance. Thanks to Rodolfo Dirzo for providing coworking space and Tatjana Schlechtweg for manuscript comments. Thanks to Vladimir Kirilin for assistance with specimen collection. Thanks to Dmitri Petrov and Bernard Kim for feedback on the project. Thanks to Liz Hadly for mentorship and use of lab space. Funding was provided by the National Science Foundation Postdoctoral Research Fellowship in Biology, the PRISM Baker Fellowship, the Howard Hughes Medical Institute, National Science Foundation grant 2209394, and the Irene Brown Fund for Nature-Related Research and Environmental Justice. The work of E.Y. was conducted under the IDB RAS Government basic research program in 2026 No. 0088-2024-0017. Undergraduate trainees were funded through the Biology Summer Undergraduate Research Program, Mentoring Undergraduates in Interdisciplinary Research, and Community College Outreach Program programs at Stanford University.

## Authors’ Contributions

KIV conceptualized the research project; all authors conducted investigations; KIV developed methodologies, curated data, analyzed the data, administered the project, supervised undergraduate mentees, visualized the data, and wrote the original draft; KIV, DH, EY, and BB contributed reviews and edits of the manuscripts. All authors read and approved the final manuscript.

## Statements and Declarations

The authors declare no conflict of interest.

## Declarations

### Ethics Approvals

No approval of research ethics committees was required to accomplish the goals of this study because experimental work was conducted with an unregulated invertebrate species.

### Data Availability

Sequence data are deposited in the NCBI GenBank database, and all data relevant for the study are included in the **Online Resources**.

