## Supplementary material for "Salinity is associated with habitat use and mitochondrial lineage diversity in halophilic *Ephydra* flies (Diptera: Ephydridae)": Online Resources/Supplementary Material

**Online Resource 1.** *COI* alignment utilized in our *Ephydra* spp. phylogeny.

>PV790562_Unknown_sp._Mono_Lake_CA
---------------TATTTATTTTCGGAGCTTGAGCAGGAATAGTAGGTACATCTTTAAGAATTCTAATTCGAGCAGAATTAGGTCACCCAGG
AGCATTAATTGGTGATGACCAAATTTATAATGTAATTGTTACTGCTCATGCTTTTGTAATAATTTTTTTATAGTTATACCTATTATAATT
GGAGGGTTTGGAAATTGATTAGTTCCATTAATATTAGGAGCCCCAGATATAGCATTCCCTCGAATAAATAATATAAGTTTTTGATTATT
ACCCCCAGCTTTAACTCTACTGCTAGTAAGTAGTATAGTAGAAAACGGGGCTGGTACAGGTTGAACTGTTTACCCACCACTATCAG
CAGGAATTGCTCATG--GAGGAGCATCAGTTGATTTAGCAATTTTTTCACTACATTTAGCAGGAATTTCTTCTATTCTAGGAGCCGTA
AATTTTATTACTACAGTAATTAATATACGAGCTACTGGAATTACATTTGATCGAATACCATTATTTGTATGATCAGTAGTAATTACAGCC
TTATTATTATTATTATCTTTACCTGTTTTAGCAGGTGCTATTACTATGTTATTAACAGACCGAAATTTAAATACTTCATTTTTCGATCCGG
CAGGANGGGGTGACCCAATTTTATATCAACACTTA------------------------
>PV790570_Unknown_sp._Great_Salt_Lake_UT
GATATTGGAACTCTTTATTTATTTTCGGAGCTTGAGCAGGAATAGTAGGTACATCTTTAAGAATTCTAATTCGAGCAGAATTAGGTC
ACCCAGGAGCATTAATTGGTGATGACCAAATTTATAATGTAATTGTTACTGCTCATGCTTTTGTAATAATTTTTTTATAGTTATACCTA
TTATAATTGGAGGGTTTGGAAATTGATTAGTTCCATTAATATTAGGAGCCCCAGATATAGCATTCCCTCGAATAAATAATATAAGTTTT
TGATTATTACCCCCAGCTTTAACTCTACTGCTAGTAAGTAGTATAGTAGAAAACGGGGCTGGTACAGGTTGAACTGTTTACCCACCA
CTATCAGCAGGAATTGCTCATG--GAGGAGCATCAGTTGATTTAGCAATTTTTTCACTACATTTAGCAGGAATTTCCTCTATTCTAGG
AGCCGTAAATTTTATTACTACAGTAATTAATATACGAGCTACTGGAATTACATTTGATCGAATACCATTATTTGTATGATCAGTAGTAAT
TACAGCCTTATTATTATTATTATCTTTACCTGTTTTAGCAGGTGCTATTACTATGTTATTAACAGACCGAAATTTAAATACTTCATTTTTC
GATCCGGCAGGAGGGGGTGACCCAATTTTATATCAACACTTATTTTGATTTTTTGGTCA-------
>KU496737.1_E._riparia_Alaska_
---------------------TTTTCGGAGCTTGAGCAGGAATAGTAGGTACATCTTTAAGAATTCTAATTCGAGCAGAATTAGGTCACCCAGGAG
CATTAATTGGTGATGACCAAATTTATAATGTAATTGTTACTGCTCATGCTTTTGTAATAATTTTTTTATAGTTATACCTATTATAATTGG
AGGGTTTGGAAATTGATTAGTTCCATTAATATTAGGAGCCCCAGATATAGCATTCCCTCGAATAAATAATATAAGTTTTTGATTATTAC
CCCCAGCTTTAACTCTACTGCTAGTAAGTAGTATAGTAGAAAACGGGGCTGGTACAGGTTGAACTGTTTACCCACCACTATCAGCA
GGAATTGCTCATG--GAGGAGCATCAGTTGATTTAGCAATTTTTTCACTACATTTAGCAGGAATTTCCTCTATTCTAGGAGCCGTAAA
TTTTATTACTACAGTAATTAATATACGAGCTACTGGAATTACATTTGATCGAATACCATTATTTGTATGATCAGTAGTAATTACAGCCTT
ATTATTATTATTATCTTTACCTGTTTTAGCAGGTGCTATTACTATGTTATTAACAGACCGAAATTTAAATACTTCATTTTTCGATCC--------
-----------------------------------------------------
>JF874959.1_Ephydra_sp._Manitoba_Canada_
--------AACTCTTTATTTATTTTCGGAGCTTGAGCAGGAATAGTAGGTACATCTTTAAGAATTCTAATTCGAGCAGAATTAGGTCACCC
AGGAGCATTAATTGGTGATGACCAAATTTATAATGTAATTGTTACTGCTCATGCTTTTGTAATAATTTTTTTATAGTTATACCTATTATA
ATTGGAGGGTTTGGAAATTGATTAGTTCCATTAATATTAGGAGCCCCAGATATAGCATTCCCTCGAATAAATAATATAAGTTTTTGATT
ATTACCCCCAGCTTTAACTCTACTGCTAGTAAGTAGTATAGTAGAAAACGGGGCTGGTACAGGTTGAACTGTTTACCCACCACTAT
CAGCAGGAATTGCTCATG--GAGGAGCATCAGTTGATTTAGCAATTTTTTCACTACATTTAGCAGGAATTTCCTCTATTCTAGGAGCC
GTAAATTTTATTACTACAGTAATTAATATACGAGCTACTGGAATTACATTTGATCGAATACCATTATTTGTATGATCAGTAGTAATTACA
GCCTTATTATTATTATTATCTTTACCTGTTTTAGCAGGTGCTATTACTATGTTATTAACAGACCGAAATTTAAATACTTCATTTTTCGATC
CGGCAGGAGGGGGTGACCCAATTTTATATCAACACTTATTT---------------------
>EU493562.1_E._packardi_
-------------------------------------------------------------------------------------------------------------------------------------------------------------------------------
---------------------------------------------------------ATATAGCATTCCCTCGAATAAATAATATAAGTTTTTGATTATTACCCCCAGCTTTAACTCT
ACTGCTAGTAAGTAGTATAGTAGAAAACGGGGCTGGTACAGGTTGAACTGTTTACCCACCACTATCAGCAGGAATTGCTCATG--GA
GGAGCATCAGTTGATTTAGCAATTTTTTCACTACATTTAGCAGGAATTTCCTCTATTCTAGGAGCCGTAAATTTTATTACTACAGTAAT
TAATATACGAGCTACTGGAATTACATTTGATCGAATACCATTATTTGTATGATCAGTAGTAATTACAGCCTTATTATTATTATTATCTTTA
CCTGTTTTAGCAGGTGCTATTACTATGTTATTAACAGACCGAAATTTAAATACTTCATTTTTCGATCC------------------------------------------
-------------------
>_MG081213.1_Ephydra_sp._Ontario_Canada
--------------------------------------------GTAGGTACATCTTTAAGAATTCTAATTCGAGCAGAATTAGGTCACCCAGGGGCATTAATTGGTG
ATGACCAAATTTATAATGTAATTGTTACTGCTCATGCTTTTGTAATAATTTTTTTATAGTTATACCTATTATAATTGGAGGGTTTGGAAA
TTGATTAGTTCCATTAATATTAGGAGCCCCAGATATAGCATTCCCTCGAATAAATAATATAAGTTTTTGATTATTACCCCCAGCTTTAAC
TCTACTGCTAGTAAGTAGTATAGTAGAAAACGGGGCTGGTACAGGTTGAACTGTTTACCCACCACTATCAGCAGGAATTGCTCATG
--GAGGAGCATCAGTTGATTTAGCAATTTTTTCACTACATTTAGCAGGAATTTCCTCTATTCTAGGAGCCGTAAATTTTATTACTACAG
TAATTAATATACGAGCTACTGGAATTACATTTGATCGAATACCATTATTTGTATGATCAGTAGTAATTACAGCCTTATTATTATTATTATC
TTTACCTGTTTTAGCAGGTGCTATTACTATGTTATTAACAGAC------------------------------------------------------------------------------------------
>JF867574.1_E._riparia_BOLD_
---------------------TTTTCGGAGCTTGAGCAGGAATAGTAGGTACATCTTTAAGAATTCTAATTCGAGCAGAATTAGGTCACCCAGGAG
CATTAATTGGTGATGACCAAATTTATAATGTAATTGTTACTGCTCATGCTTTTGTAATAATTTTTTTATAGTTATACCTATTATAATTGG
AGGGTTTGGAAATTGATTAGTTCCATTAATATTAGGAGCCCCAGATATAGCATTCCCTCGAATAAATAATATAAGTTTTTGATTATTAC
CCCCAGCTTTAACTCTACTGCTAGTAAGTAGTATAGTAGAAACCGGGGCTGGTACAGGTTGAACTGTTTACCCACCACTATCAGCA
GGAATTGCTCATG--GAGGAGCATCAGTTGATTTAGCAATTTTTTCACTACATTTAGCAGGAATTTCCTCTATTCTAGGAGCCGTAAA
TTTTATTACTACAGTAATTAATATACGAGCTACTGGAATTACATTTGATCGAATACCATTATTTGTATGATCAGTAGTAATTACAGCCTT
ATTATTATTATTATCTTTACCTGTTTTAGCAGGTGCTATTACTATGTTATTAACAGACCGAAATTTAAATACTTCATTTTTCGATCC--------
-----------------------------------------------------
>MF059325.1_E._macellaria_Malta_
--------AACTCTTTATTTATTTTCGGAGCTTGAGCAGGAATAGTAGGTACATCTTTAAGAATTCTAATTCGAGCAGAATTAGGTCACCC
AGGAGCATTAATTGGTGATGACCAAATTTATAATGTAATTGTTACTGCTCATGCTTTTGTAATAATTTTTTTATAGTTATACCTATTATA
ATTGGAGGGTTTGGAAATTGATTAGTTCCATTAATATTAGGAGCCCCAGATATAGCATTCCCTCGAATAAATAATATAAGTTTTTGATT
ATTACCCCCAGCTTTAACTTTATTGCTAGTAAGTAGTATAGTAGAAAACGGGGCTGGTACAGGTTGAACTGTTTACCCACCACTATC
AGCAGGAATTGCTCATG--GAGGAGCATCAGTTGATTTAGCAATTTTTTCACTACATTTAGCAGGAATTTCATCTATTCTAGGAGCCG

TAAATTTTATTACTACAGTAATTAATATACGAGCTACTGGAATTACATTTGATCGAATACCATTATTTGTATGATCAGTAGTAATTACAGC
CTTATTATTATTATTATCTTTACCTGTTTTAGCAGGTGCTATTACTATGTTATTAACAGATCGAAATTTAAATACTTCATTTTTCGATCCA
GCAGGAGGAGGTGATCCAATTTTATACCAACACTTATTT----------------------------
>MZ623564.1_E._macellaria_Finland_
----------------------------------------------------------------------------------------------------------------------------------------------------------------------------------------------------------------------------------
--------------------------------------------------------------------------ATATAGCATTCCCTCGAATAAATAATATAAGTTTTTGATTATTACCCCCAGCTTTAACTCT
ACTGCTAGTAAGTAGTATAGTAGAAAACGGGGCTGGTACAGGTTGAACTGTTTACCCACCACTATCAGCAGGAATTGCTCATG--GA
GGAGCATCAGTTGATTTAGCAATTTTTTCACTTCATTTAGCAGGAATTTCCTCTATTCTAGGGGCCGTAAATTTTATTACTACAGTAAT
TAATATACGAGCTACTGGAATTACATTTGATCGAATACCATTATTTGTATGATCAGTAGTAATTACAGCCTTATTATTATTATTATCTTTA
CCTGTTTTAGCAGGTGCTATTACTATGTTATTAACAGATCGAAATTTAAATACTTCATTTTTTGATCC--------------------------------------------------------
-----------------------
>PV790571.1_Riparia-like_Ephydra_sp._Great_Salt_Lake_UT
----------------------------------------------------------------------------------------------------------------------------------------------------------------------------------------------------------------------------------
----------------------------------------------------------------GCTCCYGANATGGCTTTYCCTCGAATAAATAATATAAGTTTTTGATTATTACCCCCAGCTTTA
ACTCTACTGCTAGTAAGTAGTATAGTAGAAAACGGGGCTGGTACAGGTTGAACTGTTTACCCACCACTATCAGCAGGAATTGCTCA
TG--GAGGAGCATCAGTTGATTTAGCAATTTTTTCACTACATTTAGCAGGAATTTCCTCTATTCTAGGAGCCGTAAATTTTATTACTAC
AGTAATTAATATACGAGCTACTGGAATTACATTTGATCGAATACCATTATTTGTATGATCAGTAGTAATTACAGCCTTATTATTATTATTA
TCTTTACCTGTTTTAGCAGGTGCTATTACTATGTTATTAACAGACCGAAATTTAAATACTTCATTTTTCGATCCGGCAGGAGGGGGT
GACCCAATTTTATATCAACACTTATTTTGATTCTTCGGMCA-------
>PQ772809_E._riparia_Russia
----------------------------------------------------------------------------------------------------------------------------------------------------------------------------------------------------------------------------------
----------------------------------------------------------------------------ATGGCNTTYCCTCGAATAAATAATATAAGTTTTTGATTATTACCCCCAGCTTTAACTCT
ACTGCTAGTAAGTAGTATAGTAGAAAACGGGGCTGGTACAGGTTGAACTGTTTACCCACCACTATCAGCAGGAATTGCTCATG--GA
GGAGCATCAGTTGATTTAGCAATTTTTTCACTACATTTAGCAGGAATTTCCTCTATTCTAGGAGCCGTAAATTTTATTACTACAGTAAT
TAATATACGAGCTACTGGAATTACATTTGATCGAATACCATTATTTGTATGATCAGTAGTAATTACAGCCTTATTATTATTATTATCTTTA
CCTGTTTTAGCAGGTGCTATTACTATGTTATTAACAGACCGAAATTTAAATACTTCATTTTTCGATCCGGCAGGNNGGGGTGACCCA
ATTTTATATCAACACTTATTTTGRTTCTTCGGNCCACCC---
>PV790564.1_E._millbrae_South_Bay_Salt_Ponds_CA
----------------------------------------------------------------------------------------------------------------------------------------------------------------------------------------------------------------------------------
----------------------------------------------------------------------------ATGGCATTCCCTCGAATAAATAATATAAGTTTTTGATTACTACCCCCAGCTTTAACTCT
TTTACTAGTAAGTAGTATAGTAGAAAACGGGGCTGGTACAGGCTGAACTGTTTACCCACCACTATCAGCAGGAATTGCTCATG--GT
GGAGCATCAGTTGATCTAGCAATTTTTTCTTTACACTTAGCAGGAATTTCTTCTATTCTAGGAGCCGTAAATTTTATTACTACAGTAAT
TAATATACGAGCCACTGGAATTACATTTGACCGAATACCATTATTTGTGTGATCAGTAGTAATTACAGCTTTATTATTATTATTATCATTA
CCTGTTTTAGCAGGAGCTATTACTATATTATTAACTGATCGAAA--------------------------------------------------------------------------------------------------------------
>PV790567_E._millbrae_South_Bay_Salt_Ponds_CA
----------------------------------------------------------------------------------------------------------------------------------------------------------------------------------------------------------------------------------
-------------------------------------------------------------------------------------------------------AATATAAGTTTTTGATTACTACCCCCAGCTTTAACTCTTTTACTAGTAA
GTAGTATAGTAGAAAACGGGGCTGGTACAGGCTGAACTGTTTACCCACCACTATCAGCAGGAATTGCTCATG--GTGGAGCATCAG
TTGATCTAGCAATTTTTTCTTTACACTTAGCAGGAATTTCTTCTATTCTAGGAGCCGTAAATTTTATTACTACAGTAATTAATATACGAG
CCACTGGAATTACATTTGACCGAATACCATTATTTGTATGATCAGTAGTAATTACAGCTTTATTATTATTATTATCATTACCTGTTTTAGC
AGGAGCTATTACTATATTATTAACTGATCGAAATTTAAATACTTCATTTTTCGACCCTGCAGGGGGAGGAGATCCAATTTTATACCAA
CACTTATT-----------------------------
>PQ772810.1_E._millbrae_Hayward_CA
----------------------------------------------------------------------------------------------------------------------------------------------------------------------------------------------------------------------------------
----------------------------------------------------------------GCWCCMGAYATGGCNTTCCCTCGAATAAATAATATAAGTTTTTGATTACTACCCCCAGCTTT
AACTCTTTTACTAGTAAGTAGTATAGTAGAAAACGGGGCTGGTACAGGCTGAACTGTTTACCCACCACTATCAGCAGGAATTGCTC
ATG--GTGGAGCATCAGTTGATCTAGCAATTTTTTCTTTACACTTAGCAGGAATTTCTTCTATTCTAGGAGCCGTAAATTTTATTACTA
CAGTAATTAATATACGAGCCACTGGAATTACATTTGACCGAATACCATTATTTGTATGATCAGTAGTAATTACAGCTTTATTATTATTATT
ATCATTACCTGTTTTAGCAGGAGCTATTACTATATTATTAACTGATCGAAATTTAAATACTTCATTTTTCGACCCTGCAGGGGGAGGA
GATCCAATTTTATACCAACACTTATTTTGRTTCTTCGGNCACCCMGAA
>PQ772806.1_E._millbrae_South_Bay_Salt_Ponds_CA
----------------------------------------------------------------------------------------------------------------------------------------------------------------------------------------------------------------------------------
--------------------------------------------------------------------------------GCWTTYCCTCGAATAAATAATATAAGTTTTTGATTACTACCCCCAGCTTTAACTCTT
TTACTAGTAAGTAGTATAGTAGAAAACGGGGCTGGTACAGGCTGAACTGTTTACCCACCACTATCAGCAGGAATTGCTCATG--GTG
GAGCATCAGTTGATCTAGCAATTTTTTCTTTACACTTAGCAGGAATTTCTTCTATTCTAGGAGCCGTAAATTTTATTACTACAGTAATT
AATATACGAGCCACTGGAATTACATTTGACCGAATACCATTATTTGTGTGATCAGTAGTAATTACAGCTTTATTATTATTATTATCATTA
CCTGTTTTAGCAGGAGCTATTACTATATTATTAACTGATCGAAATTTAAATACTTCATTTTTCGACCCTGCAGGGGGAGGAGATCCAA
TTTTATACCAACACTTATTTTGRTTCTTCGGNCACCCMGAA
>OP971562.1_E._gracilis_Great_Salt_Lake_UT
----------------------------------------------------------------------------------------------------------------------------------------------------------------------------------------------------------------------------------
--------------------------------------------------------------------------ATATAGCATTCCCTCGAATAAATAATATAAGTTTTTGATTATTACCTCCTGCTTTAACTCT
TTTACTAGTAAGTAGTATAGTAAAAAACGGGGCTGGTACAGGATGAACTGTTTACCCACCACTATCAGCAGGAATTGCTCATG--GA
GGAGCTTCAGTTGATTTAGCTATTTTTTCTCTACATTTAGCAGGTATTTCATCTATTTTAGGAGCCGTAAATTTTATTACAACTGTAAT
TAATATACGAGCTACTGGAATTACATTTGATCGAATACCATTATTTGTTTGATCAGTAGTAATTACAGCTTTATTATTATTATTATCATTA
CCTGTATTAGCAGGAGCTATTACTATATTATTAACAGATCGAAACTTAAATACATCATTTTTCGATCC--------------------------------------------------------
-----------------------
>OP971557.1_E._gracilis_Great_Salt_Lake_UT
-------------------------------------------------------------GGAACTTCATTAAGTATTTTAATTCGAGCAGAATTAGGTCATCCAGGAGCATTAATTGGAGAT
GACCAAATTTATAATGTAATTGTTACTGCTCATGCTTTTGTAATAATTTTTTTTATAGTTATACCTATTATAATTGGAGGGTTTGGAAATT
GATTAGTTCCATTAATATTAGGAGCTCCAGATATAGCATTCCCTCGAATAAATAATATAAGTTTTTGATTATTACCTCCCGCTTTAACTC

TTTTACTAGTAAGTAGTATAGTAGAAAACGGGGCTGGTACAGGATGAACTGTTTACCCACCACTATCAGCAGGAATTGCTCATG--G
AGGAGCTTCAGTTGATTTAGCTATTTTTTCTCTACATTTAGCAGGAATTTCATCTATTTTAGGAGCCGTAAATTTTATTACAACTGTAA
TTAATATACGAGCTACTGGAATTACATTTGATCGAATACCATTATTTGTTTGATCAGTAGTAATTACAGCTTTATTATTATTATTATCATT
ACCTGTATTAGCAGGAGCTATTACTATATTATTAACAGATCGAAACTTAAATACATCATTTTTCGATCCTGCAGGAGGGGGAGACCC
A-------------------------------------------
>JF867420.1_Ephydridae_sp.
--------AACTCTTTATTTTATCTTCGGAGCTTGAGCAGGAATAGTAGGAACATCTCTTAGTATTTTAATTCGAGCCGAATTAGGTCATCC
TGGAGCATTAATTGGTGATGATCAAATTTATAACGTAATTGTTACTGCTCATGCATTTGTAATAATTTTCTTTATAGTAATACCTATTATA
ATTGGAGGATTTGGAAATTGATTAGTACCTTTAATATTAGGAGCACCTGATATAGCATTTCCTCGAATAAATAATATAAGTTTTTGATTA
TTACCTCCAGCTTTAACTCTTTTACTAGTAAGTAGTATAGTAGAAAACGGAGCTGGTACTGGATGAACTGTTTACCCACCCCTATCA
GCAGGAATTGCTCATG--GAGGAGCATCAGTTGATTTAGCTATTTTTTCTCTTCATTTAGCAGGAATTTCATCTATTTTAGGAGCAGTA
AATTTTATTACAACAGTAATTAATATGCGAGCTACTGGGATTACATTTGATCGAATACCTTTATTTGTTTGATCAGTAGTTATTACAGCT
TTATTATTACTTCTATCACTACCTGTATTAGCAGGAGCAATTACTATACTATTAACAGATCGAAATTTAAATACTTCATTTTTTGACCCT
GCAGGAGGGGGAGACCCAATTTTATACCAACATTTATTT---------------------
>PQ772805.1_E._gracilis_South_Bay_Salt_Ponds_CA
---------------------------------------------------------------------------------------------------------------------------------------------------------------------------------------------
-------------------------------------------------------------------GCTTTCCCTCGAATAAATAATATAAGTTTTTGATTACTTCCTCCCGCACTAACACTT
TTACTAGTTAGTAGAATAGTAGAAAACGGAGCTGGTACAGGATGAACTGTTTACCCTCCGCTATCAGCAGGAATTGCTCATG--GAG
GTGCTTCTGTAGATATAACTATTTTTTCTCTTCATTTAGCCGGTATTTCATCTATTTTAGGAGCTGTAAATTTTATTACAACTGTAATTA
ATATACGAGCTACTGGAATTACATTCGATCGAATACCATTATTTGTTTGATCAGTAGTAATTACTGCTTTATTACTACTACTATCATTAC
CTGTATTAGCTGGAGCTATTACAATATTATTAACAGATCGAAACTTAAATACATCATTTTTTGATCCTGCAGGAGGGGGAGACCCAAT
TTTATACCAACATTTATTTTGRTTCTTCGGNCACCCMGAA
>PQ772812.1_E._gracilis_South_Bay_Salt_Ponds_CA
---------------------------------------------------------------------------------------------------------------------------------------------------------------------------------------------
-----------------------------------------------------------------------------------------NNNNNNNNNTNNNTTANTNCTNCCGCACTAACACTTTTACTAGTT
AGTAGAATAGTAGAAAACGGAGCTGGTACAGGATGAACTGTTTACCCTCCGCTATCAGCAGGAATTGCTCATG--GAGGTGCTTCT
GTAGATATAACTATTTTTTCTCTTCATTTAGCCGGTATTTCATCTATTTTAGGAGCTGTAAATTTTATTACAACTGTAATTAATATACGAG
CTACTGGAATTACATTCGATCGAATACCATTATTTGTTTGATCAGTAGTAATTACTGCTTTATTACTACTACTATCATTACCTGTATTAG
CTGGAGCTATTACAATATTATTAACAGATCGAAACTTAAATACATCATTTTTTGATCCTGCAGGAGGGGGAGACCCAATTTTATACCA
ACATTTATTTTGATTCTTCGGNCACCCCGAA
>PV790568.1_E._gracilis_South_Bay_Salt_Ponds_CA
---------------------------------------------------------------------------------------------------------------------------------------------------------------------------------------------
------------------------------------------------------GCTCCTNNNATGGCNTTCCCTNGAATAAATAATATAAGTTTTTGATTACTTCCTCCCGCACTA
ACACTTTTACTAGTTAGTAGAATAGTAGAAAACGGAGCTGGTACAGGATGAACTGTTTACCCTCCGCTATCAGCAGGAATTGCTCA
TG--GAGGTGCTTCTGTAGATATAACTATTTTTTCTCTTCATTTAGCTGGTATTTCATCTATTTTAGGAGCTGTAAATTTTATTACAACT
GTAATTAATATACGAGCTACTGGAATTACATTCGACCGAATACCATTATTTGTTTGATCAGTAGTAATTACTGCTTTATTATTACTACTA
TCATTACCTGTATTAGCTGGAGCTATTACAATATTATTAACAGATCGAAACTTAAATACATCNNNNNNNNNNNNNNNN-------------------
-----------------------------------------
>PQ772817.1_E._gracilis_South_Bay_Salt_Ponds_CA
---------------------------------------------------------------------------------------------------------------------------------------------------------------------------------------------
------------------------------------------------------------------------TTCCCTAAAAAAAAAAAATAAGTTTTTGATTACTTCCTCCTGCATTAACACTTTTA
CTAGTTAGTAGAATAGTAGAAAACGGAGCTGGTACAGGATGAACTCTTTACCCTCCGCTATCAGCAGGAATTGCACATG--GAGGT
GCTTCTGTAGATTTAACTATTTTTTCTTTTCATTTAGCCGGTATTTCATCAATTTTAGGAGATGTAAATTTTATTACAACTGTAATTAATA
TACGAGCTACTGGAATTACATTTGACCGAATACCATTATTTGTATGATCAGTAGTAATTACTGCTTTATTATTATTACTATCATTACCTG
TATTAGCTGGAGCTATTACAATATTATTAACAGATCGAAATTTAAATACATCATTTTTT-------------------------------------------------------------------
----
>PV790563.1_E._hians_Lake_Abert_OR
---------------------------------------------------------------------------------------------------------------------------------------------------------------------------------------------
----------------------------------------------------------------ATAGCCTTTCCTCGAATAAATAATATAAGTTTTTGATTATTACCTCCTGCTTTAACTCTT
TTACTAGTAAGTAGCATAGTAGAAAATGGGGCTGGTACAGGTTGAACTGTTTACCCACCTTTATCCGCAGGAATTGCTCATG--GAG
GAGCTTCAGTTGACTTAGCTATTTTCTCTCTTCATTTAGCTGGAATTTCTTCAATTTTAGGGGCCGTAAATTTTATTACTACTGTAATT
AATATACGAGCTACTGGAATTACATTTGATCGAATACCTTTATTCGTTTGATCAGTAGTTATTACAGCTCTTTTATTATTATTATCACTA
CCAGTATTAGCAGGAGCAATTACTATACTTCTTACTGACCGAAATTTAAATACATCCTTTTTTGACCCAGCTG--------------------------------
----------------------------
>OP971547.1_E._hians_Great_Salt_Lake_UT
---------------------------------------------------GGAACATCATTAAGAATTCTAATTCGAGCAGAATTAGGACATCCTGGGGCATTAATTGGTGAT
GACCAAATTTATAATGTAATTGTTACTGCCCATGCTTTTGTAATAATTTTCTTCATAGTAATACCTATTATAATTGGAGGATTTGGAAAT
TGATTAGTTCCTTTAATATTAGGAGCTCCTGATATAGCCTTTCCTCGAATAAATAATATAAGTTTTTGATTATTACCTCCTGCTTTAACT
CTTTTACTAGTAAGTAGCATAGTAGAAAATGGGGCTGGTACAGGTTGAACTGTTTACCCACCTTTATCCGCAGGAATTGCTCATG--
GAGGAGCTTCAGTTGACTTAGCTATTTTCTCTCTTCATTTAGCTGGAATTTCTTCAATTTTAGGAGCCGTAAATTTTATTACTACTGT
AATTAATATACGAGCTACTGGAATTACATTTGATCGAATACCTTTATTCGTTTGATCAGTAGTTATTACAGCTCTTTTATTATTATTATCA
CTACCAGTATTAGCAGGAGCAATTACTATACTTCTTACTGACCGAAATTTAAATACATCCTTTTTTGATCCAGCTGGAGGAGGAGAC
CCT---------------------------------------------
>HM374264.1_E._hians_BOLD
--------TACTCTTTATTTTATTTTTGGAGCATGAGCAGGAATAGTGGGAACATCATTAAGAATTCTAATTCGAGCAGAATTAGGACATCC
TGGAGCATTAATTGGTGATGATCAAATTTATAATGTAATTGTTACTGCCCATGCTTTTGTAATAATTTTCTTCATAGTAATACCTATTATA
ATTGGAGGATTTGGAAATTGATTAGTTCCTTTAATATTAGGAGCTCCTGATATAGCCTTTCCTCGAATAAATAATATAAGTTTTTGATTA
TTACCTCCTGCTTTAACTCTTTTACTAGTAAGTAGCATAGTAGAAAATGGGGCTGGTACAGGTTGAACTGTTTACCCACCTTTATCC
GCAGGGATTGCTCATG--GAGGAGCTTCAGTTGACTTAGCTATTTTCTCTCTTCATTTAGCTGGAATTTCTTCAATTTTAGGAGCCGT
AAATTTTATTACTACTGTAATTAATATACGAGCTACTGGAATTACATTTGATCGAATACCTTTATTCGTTTGATCAGTAGTTATTACAGC

TCTTTTATTATTATTATCACTACCAGTATTAGCAGGAGCAATTACTATACTTCTTACTGACCGAAATTTAAATACATCCTTTTTTGATCC
AGCTGGAGGAGGAGACCCTATTTTATACCAACATTTATTT----------------------
>PQ772816.1_E._hians_Lee_Vining_CA
--------------------------------------------------------------------------------------------------------------------------------------------------------------------------------
---------------------------------------------------------ATATAGCCTTTCCTCGAATAAATAATATAAGTTTTTGATTATTACCTCCTGCTTTAACTCT
TTTACTAGTAAGTAGTATAGTAGAAAACGGGGCTGGTACAGGTTGAACTGTTTACCCACCTTTATCCGCAGGGATTGCTCATG--GA
GGGGCTTCAGTTGACTTAGCTATTTTCTCTCTCCATTTAGCTGGAATTTCTTCAATTTTAGGAGCCGTAAATTTTATTACTACTGTAA
TTAATATACGAGCTACTGGAATTACATTTGATCGAATACCTTTATTCGTTTGATCAGTAGTTATTACAGCTCTTTTATTATTATTATCACT
ACCAGTATTAGCAGGAGCAATTACTATACTTCTTACTGACCGAAATTTAAATACATCCTTTTTTGATCCAGCTG-----------------------------
---------------------------
>PV790569_E._hians_Great_Salt_Lake_UT
--------------------------------------------------------------------------------------------------------------------------------------------------------------------------------
--------------------------------------------------------------GCTTTTCCTCGAATAAATAATATAAGTTTTTGATTATTACCTCCTGCTTTAACTCTTT
TACTAGTAAGTAGTATAGTAGAAAACGGGGCTGGTACAGGTTGAACTGTTTACCCACCTTTATCTGCAGGGATTGCTCATG--GAGG
GGCTTCAGTTGACTTAGCTATTTTTTCTCTTCATTTAGCTGGAATTTCTTCAATTTTAGGAGCCGTAAATTTTATTACTACTGTAATTA
ATATACGAGCTACTGGAATTACATTTGATCGAATACCTTTATTCGTTTGGTCAGTAGTTATTACAGCTCTTTTATTATTATTATCACTAC
CAGTATTAGCAGGAGCAATTACTATACTTCTTACTGACCGAAATTTAAATACATCCTTTTTTGACCCAGCTGGAGGAGGAGATCCTA
TTTTATACCAACATTTATTTTGATTCTTCGGMCA-------
>PV790572_E._hians_Lake_Abert_OR
--------------------------------------------------------------------------------------------------------------------------------------------------------------------------------
--------------------------------------------------------------------CCTCGAATAAATAATATAAGTTTTTGATTATTACCTCCTGCTTTAACTCTTTTAC
TAGTAAGTAGTATAGTAGAAAACGGGGCTGGTACAGGTTGAACTGTTTACCCACCTTTATCTGCAGGGATTGCTCATG--GAGGGG
CTTCAGTTGACTTAGCTATTTTCTCTCTTCATTTAGCTGGAATTTCTTCAATTTTAGGAGCCGTAAATTTTATTACTACTGTAATTAATA
TACGAGCTACTGGAATTACATTTGATCGAATACCTTTATTCGTTTGATCAGTAGTTATTACAGCTCTTTTATTATTATTATCACTACCAG
TATTAGCAGGAGCAATTACTATACTTCTTACTGACCGAAATTTAAATACATCCTTTTTTGACCCAGCTGGAGGAGGAGATCCTATTTT
ATACCAACATTTA------------------------
>PV790573.1_E._hians_Mono_Lake_CA
GATATTGGTACTCTTTATTTTATTTTTGGAGCATGAGCAGGAATAGTAGGAACATCTTTAAGAATTCTAATTCGAGCAGAATTAGGAC
ATCCTGGAGCGTTAATTGGTGATGACCAAATTTATAATGTAATTGTTACTGCCCATGCTTTTGTAATAATTTTCTTCATAGTAATACCT
ATTATAATTGGAGGATTTGGAAATTGATTAGTTCCTTTAATACTAGGAGCTCCTGATATAGCCTTTCCTCGAATAAATAATATAAGTTTT
TGATTATTACCTCCTGCTTTAACTCTTTTACTAGTAAGTAGTATAGTAGAAAACGGGGCTGGTACAGGTTGAACTGTTTACCCACCT
TTATCTGCAGGGATTGCTCATG--GAGGGGCTTCAGTTGACTTAGCTATTTTCTCTCTTCATTTAGCTGGAATTTCTTCAATTTTAGG
AGCCGTAAATTTTATTACTACTGTAATTAATATACGAGCTACTGGAATTACATTTGATCGAATACCTTTATTCGTTTGATCAGTAGTTAT
TACAGCTCTTTTATTATTATTATCACTACCAGTATTAGCAGGAGCAATTACTATACTTCTTACTGACCGAAATTTAAATACATCCTTTTT
TGACCCAGCTGGAGGAGGAGATCCTATTTTATACCAACATTTATTTTGATTTTTTGGTCA-------
>PQ772813.1_E._hians_Great_Salt_Lake_UT
--------------------------------------------------------------------------------------------------------------------------------------------------------------------------------
--------------------------------------------------------------GCTTTTCCTCGAATAAATAATATAAGTTTTTGATTATTACCTCCTGCTTTAACTCTTT
TACTAGTAAGTAGTATAGTAGAAAACGGGGCTGGTACAGGTTGAACTGTTTACCCACCTTTATCCGCAGGGATTGCTCATG--GAGG
GGCTTCAGTTGACTTAGCTATTTTCTCTCTCCATTTAGCTGGAATTTCTTCAATTTTAGGAGCCGTAAATTTTATTACTACTGTAATTA
ATATACGAGCTACTGGAATTACATTTGATCGAATACCTTTATTCGTTTGATCAGTAGTTATTACAGCTCTTTTATTATTATTATCACTAC
CAGTATTAGCAGGAGCAATTACTATACTTCTTACTGACCGAAATTTAAATACATCCTTTTTTGATCCAGCTGGAGGGGGAGACCCTA
TTTTATACCAACATTTATTTTGATTCTTCGGMCA-------
>PQ772814.1_E._hians_Great_Salt_Lake_UT
--------------------------------------------------------------------------------------------------------------------------------------------------------------------------------
--------------------------------------------------------------GCTTTTCCTCGAATAAATAATATAAGTTTTTGATTATTACCTCCTGCTTTAACTCTTT
TACTAGTAAGTAGCATAGTAGAAAATGGGGCTGGTACAGGTTGAACTGTTTACCCACCTTTATCCGCAGGAATTGCTCATG--GAGG
AGCTTCAGTTGACTTAGCTATTTTCTCTCTTCATTTAGCTGGAATTTCTTCAATTTTAGGGGCCGTAAATTTTATTACTACTGTAATTA
ATATACGAGCTACTGGAATTACATTTGATCGAATACCTTTATTCGTTTGATCAGTAGTTATTACAGCTCTTTTATTATTATTATCACTAC
CAGTATTAGCAGGAGCAATTACTATACTTCTTACTGACCGAAATTTAAATACATCCTTTTTTGATCCAGCTGGAGGAGGAGACCCTA
TTTTATACCAACATTTATTTTGATTCTTCGGMCA-------
>PQ772808.1_E._hians_Deep_Springs_CA
--------------------------------------------------------------------------------------------------------------------------------------------------------------------------------
----------------------------------------------------------------------------------------------------------------ANCTCTTTTACTAGTAAGTAGCATAGTAGAAA
ATGGGGCTGGTACAGGTTGAACTGTTTACCCACCTTTATCCGCAGGGATCGCTCATG--GAGGAGCTTCAGTTGACTTAACTATTTT
CTCTCTTCATTTAGCTGGAATTTCTTCAATTTTAGGAGCCGTAAATTTTATTACTACTGTAATTAATATACGAGCTACTGGAATTACATT
TGATCGAATACCTTTATTCGTTTGGTCAGTAGTTATTACAGCTCTTTTATTATTATTATCACTACCAGTATTAGCAGGAGCAATTACTAT
ACTTCTTACTGACCGAAATTTAAATACATCCTTTTTTGACCCAGCTGGAGGAGGAGACCCTATTTTATATCAACATTTATT-----------------
-------
>PQ772815.1_E._hians_Owens_Lake_CA
--------------------------------------------------------------------------------------------------------------------------------------------------------------------------------
---------------------------------------------------------ATATAGCCTTTCCTCSAAWWAATAATATAAGTTTTKSATTATKACCTCCTGCTTTAACTC
TTTTACTAGTAAGTAGTATAGTAGAAAAKGGGGCTGGTACAGGTTGAACTGTTTACCCACCTTTATMCGCAGGAATTGCTCATG--G
AGGAGCTTCAGTTGMYTTAGCTATTTTCTYTCTTCATTTAGCTGGAATTTCTTCAATTTTAGGGGCCGTAAAWTTTATTACTACTGTA
ATTAATATACGAGCTACTGGAATTACATTTGATCGAATACCTTTATTCGTTTGATCAGTAGTTATTACAGCTCTTTTATTATTATTATCAC
TACCAGTATTAGCASGAGCAATTACTATACTTCTTACTGACCGAAATTTAAATACATCCTTTTTTGACCC--------------------------------------
-----------------------
>PQ772807.1_E._hians_Big_Soda_Lake_CO
--------------------------------------------------------------------------------------------------------------------------------------------------------------------------------
---------------------------------------------------------ATATARCCTTTCCTCKAATAAATAATATAAGTTTTTKATTWTTACYTCYTGCTTTAACTCT

TTTACTAGTAAGDAGTATAGTAGAAAACGGGGSTGGTACAGGTTGAACTGTTTACCCMCCTYTATCCGCAGGGATTGCTCATG--GA
GGDGCTTCAGTTGAYTTAGCTATTTTCTYTCTYCATTTAGCTGGAATTTCTTCAATTTTAGGAGCCGTAAATTTTATTACTACTGTAAT
TAATATACGAGSTACTGGAATTWCATTTGATYGAATASCWTTATTYNYWTGAYCNSTAKTTATTWCANCTCTTYTATTATTATTATCAC
TACCARTATTANYRSRAGCNRTYAYKATACTWCTTACTGACCKWNATTTAMMTACATCMTTKTTWGATCCNACTG---------------------
--------------------------------
>PV790574.1_Unknown_sp._Walker_Lake,_Nevada
-----------------------------GCATGAGCAGGAATAGTAGGAACTTCTTTAAGAATTTTAATTCGAGCAGAATTAGGACATCCAGGAGCTTTA
ATTGGAGATGATCAAATTTATAATGTTATTGTAACTGCACACGCTTTTGTAATAATTTTTTTTATAGTAATACCTATTATAATTGGAGGAT
TTGGAAATTGACTAGTTCCATTAATATTAGGAGCTCCAGATATAGCATTCCCTCGAATAAATAATATAAGTTTTTGATTATTACCCCCA
GCTCTGACTCTTTTACTAGTAAGTAGTATAGTAGAAAATGGATCTGGTACAGGATGAACTGTTTACCCACCTCTATCAGCTGGTATT
GCTCATG--GAGGAGCTTCTGTTGATTTAACTATTTTTTCTCTTCATTTAGCAGGAATTTCTTCTATTTTAGGAGCTGTAAATTTTATTA
CAACTGTAATTAATATACGAGCAACAGGTATTACTTTTGATCGAATACCTTTATTTGTATGATCAGTTGTAATTACAGCTTTACTTTTAT
TATTATCTTTACCAGTTTTAGCAGGAGCTATTACAATACTATTAACAGATCGAAATTTAAATACTTCTTTTTTCGACCCA-----------------
----------------------------------------
>PZ634866.1_Riparia-like_sp_GSL_UT
----------AAAGATATTGGAACTTCGGAGTTTG---AGGAATAGTAGGAACATCATTAAGAATTTTAATTCGAGCAGAATTAGGTCACCCA
GGTGCATTAATTGGTGATGCTCAAATTTTTAATGTCATTGTTACTACTCATGCTTTTGTAATAATTTTTTTTAAAGGAATACCTATTGTA
ATTGGAGGATTTAGAACGGGATTAGTTCTATTGATACTYGGAGCACCAGATATAGTATTCCCTCGAATAAATAATATAAGTTTTTGATT
ACTACCTCCTGCATTAACTCTATTGCTAGTAAGTAGTATAGTAAAAAACGGGGCTGGCACAGGTTGAACTGTATGCCCACCACTAT
CAACAGRAATTGCACATG--GTGGAGCATTAGTTGATTTAGCAATTTTTTCAGTTCATTTAGCAGGAATTTCTTCTATTTTAGGAGCA
GTAAATTTTATCACTACAGTAATTAATATACGAGCTACTGGAATTACATTTGATCGAATACCATTATTTATATGATTAGTAGTTATTACTG
CTTGATTATTACCATTATCATTACCTGTTTTAGCTGGAGCTATTACTATGTTACTATCAGAACGAAATTTCAATACTTAA------------------
---------------------------------------------------
>AB669723.1_Drosophila_punjabensis_
GATATTGGAACTTTATATTTTATTTTTGGAGCTTGAGCAGGAATAGTAGGAACATCTTTAAGAATTTTAATTCGAGCTGAATTAGGAC
ATCCTGGAGCTTTAATTGGAGATGATCAAATTTATAATGTAATTGTAACAGCTCATGCTTTTATTATAATTTTTTTTATAGTTATACCTAT
TATAATTGGAGGATTTGGTAATTGATTAGTTCCTTTAATATTAGGAGCTCCTGATATAGCATTCCCTCGAATAAATAATATAAGATTTTG
ATTATTACCTCCTGCTCTTTCACTTTTATTAGTAAGCAGAATGGTTGAAAATGGAGCTGGTACTGGATGAACAGTTTATCCACCGTT
ATCTTCAGGAATTGCCCATG--GAGGAGCTTCTGTAGATTTAGCTATTTTTTCTTTACATTTAGCTGGAATTTCATCAATTTTAGGAGC
TGTAAATTTTATTACTACAGTAATTAATATACGATCTACAGGAATTACATTAGATCGAATACCATTATTTGTATGATCAGTTGTAATTACT
GCTTTATTACTTTTATTATCATTACCAGTACTAGCTGGAGCAATTACAATATTACTAACAGATCGAAATTTAAATACATCATTTTTCGAC
CCTGCTGGAGGTGGAGATCCAATTTTATATCAACATTTATTTTGATTTTTTGGACA-------
>KU875234.1_Hydrellia_griseola_
--------AACCCTATATTTTATTTTCGGTGCATGATCCGGAATAGTTGGAACTTCATTAAGTATTTTAATTCGAGCTGAACTGGGTCACCC
GGGAGCTTTAATTGGTGATGACCAAATTTATAACGTAATTGTTACAGCTCATGCATTTATTATAATTTTCTTTATGGTAATACCTATTAT
AATTGGAGGATTTGGAAATTGATTAGTTCCTTTAATATTAGGAGCCCCTGATATAGCCTTTCCTCGAATAAATAATATAAGTTTTTGAT
TACTACCCCCAGCTCTTACCTTATTATTAGTCAGTAGTATAGTAGATAACGGAGCTGGAACAGGATGAACGGTTTACCCTCCACTTT
CTTCTGGAATTGCTCATG--GAGGAGCTTCTGTAGATCTTGCTATTTTTTCTCTTCATTTAGCAGGAATTTCTTCAATTTTAGGAGCA
GTAAATTTTATTACTACTGTAATTAATATACGATCAACAGGTATTACATTTGACCGAATACCTTTATTTGTATGATCTGTTGTTATTACAG
CTTTATTACTACTTTTATCATTACCTGTTTTAGCCGGAGCTATTACAATATTATTAACTGACCGAAATTTAAATACATCATTTTTTGACC
CTGCTGGAGGAGGAGACCCTATTCTTTACCAACACTTATTT--------------------
>GU673025.1_Maniola_jurtina
--------AACATTGTATTTTATTTTCGGAGCTTGATCAGGAATAGTTGGAACTTCTTTAAGTATTTTAATTCGAGCTGAACTAGGACATCC
TGGAGCTCTTATTGGTGACGATCAAATTTATAACGTAATTGTAACAGCCCATGCTTTTATTATAATTTTTTTTATGGTAATACCTATTAT
AATTGGAGGATTTGGAAACTGATTAGTTCCTCTTATACTAGGTGCACCTGATATAGCTTTCCCTCGAATAAATAATATAAGTTTTTGA
CTTTTACCTCCTGCATTAACTTTATTGTTAGTAAGTAGTATAGTAGAAAACGGAGCTGGAACAGGATGAACTGTTTACCCTCCCCTA
TCTTCTAATATTGCTCATG--GCGGAGCTTCTGTTGATTTAGCTATTTTTTCTCTTCATTTAGCTGGAATTTCATCAATTTTAGGAGCA
GTAAATTTTATTACTACTGTAATTAATATACGATCAACAGGTATTACCTTTGATCGAATACCTTTATTTGTTTGATCTGTAGTAATTACAG
CTTTATTACTTTTATTATCTTTACCTGTTTTAGCTGGAGCTATTACTATATTATTAACAGATCGAAATTTAAATACATCATTTTTTGATCC
AGCAGGAGGAGGAGATCCAATTTTATACCAACATTTATTT--------------------

**Online Resource 2.** *Ephydra* spp. *COI* genealogy newick file.

```
(_PV790570__Unknown_sp.__Great_Salt_Lake__UT__:0.0000027135,_JF874959.1__Ephydra_sp.__M
anitoba__Canada__:0.0000010000,(_KU496737.1__E._riparia__Alaska__:0.0000010000,((_EU493562.1
__E._packardi_:0.0000010000,(((PV790562_Unknown_sp.__Mono_Lake__CA__:0.0000024413,(_MF05
9325.1__E._macellaria__Malta__:0.0033693395,((((_PQ772810.1__E._millbrae__Hayward__CA_:0.000
0010000,(_PQ772806.1__E._millbrae__South_Bay_Salt_Ponds__CA_:0.0000010000,_PV790564.1__E.
_millbrae__South_Bay_Salt_Ponds__CA_:0.0000010000)99:0.0023288822)66:0.0000026081,PV790567
_E._millbrae__South_Bay_Salt_Ponds__CA_:0.0000023433)98:0.0325774627,((_OP971557.1__E._grac
ilis__Great_Salt_Lake__UT_:0.0000020473,(_OP971562.1__E._gracilis__Great_Salt_Lake__UT_:0.005
7344143,(PV790568.1_E._gracilis__South_Bay_Salt_Ponds__CA_:0.0000020144,((PQ772805.1_E._gra
cilis__South_Bay_Salt_Ponds__CA_:0.0000023388,PQ772812.1_E._gracilis__South_Bay_Salt_Ponds_
_CA_:0.0051282379)93:0.0053473286,PQ772817.1_E._gracilis__South_Bay_Salt_Ponds__CA_:0.0640
523577)79:0.0023293035)100:0.0789232425)55:0.0023709897)64:0.0155491777,((JF867420.1_Ephydri
dae_sp.:0.0479625970,(((PV790573.1_E._hians__Mono_Lake__CA_:0.0000010000,PV790572_E._hian
s__Lake_Abert__OR_:0.0000010000)89:0.0075648547,(((PQ772816.1_E._hians__Lee_Vining__CA_:0.
0000010000,PQ772813.1_E._hians__Great_Salt_Lake__UT_:0.0035566007)49:0.0000025403,PQ77280
7.1_E._hians__Big_Soda_Lake__CO_:0.0028533487)56:0.0034761063,((HM374264.1_E._hians__BOL
D_:0.0000026121,PQ772808.1_E._hians__Deep_Springs__CA_:0.0142045602)56:0.0016777183,(OP97
1547.1_E._hians__Great_Salt_Lake__UT_:0.0000020300,((PV790563.1_E._hians__Lake_Abert__OR_:
0.0000010000,PQ772815.1_E._hians__Owens_Lake__CA_:0.0026975049)94:0.0026141117,PQ772814.
1_E._hians__Great_Salt_Lake__UT_:0.0023523287)88:0.0023448322)88:0.0034529477)87:0.00664251
95)51:0.0064532787)48:0.0037991370,PV790569_E._hians__Great_Salt_Lake__UT_:0.0030959397)10
0:0.0685502248)76:0.0154883621,(PV790574.1_Unknown_sp.___Walker_Lake__Nevada_:0.058037770
8,(AB669723.1_Drosophila_punjabensis_:0.0821841466,(KU875234.1_Hydrellia_griseola_Dipt_outgroup
:0.0862293322,GU673025.1_Maniola_jurtina_:0.0784693898)98:0.0380756294)94:0.0371228171)71:0.0
228011750)46:0.0136774340)97:0.0208605527)91:0.0140162438,PZ634866_Riparia-like_sp_GSL_UT:0.
1756943981)96:0.0117163190)96:0.0101981113)75:0.0015947841,(_PV790571.1__Riparia-like_Ephydra
_sp.__Great_Salt_Lake__UT_:0.0000025200,PQ772809_E._riparia__Russia_:0.0026603688)97:0.01338
71961)50:0.0028543498,(__MG081213.1__Ephydra_sp.__Ontario__Canada_:0.0000025990,_MZ62356
4.1__E._macellaria__Finland__:0.0107563977)59:0.0021866356)33:0.0000023829)43:0.0000022709,_J
F867574.1__E._riparia__BOLD__:0.0018043420)45:0.0000028642)75:0.0000015987);
```

**Online Resource 3.** Dated measurements for salinity (ppt) and pH in identified SBSPRP ponds. "NA" indicates the recorded measurement was not taken, lost or illegible.

| Date | PondName | Sal | pH |
|---|---|---|---|
| 3/28/2024 | N2 | 57 | 8.23 |
| 3/28/2024 | E3C | 13 | NA |
| 4/9/2024 | A12 | 150 | NA |
| 4/9/2024 | A11 | 15 | 8.5 |
| 4/9/2024 | A13 | 95.8 | 8.3 |
| 4/9/2024 | A14 | 116 | 7.58 |
| 4/12/2024 | S5 | 33 | NA |
| 4/12/2024 | R5 | 30 | 8.6 |
| 5/23/2024 | SF2 | NA | NA |
| 5/23/2024 | R2 | 96 | 7.14 |
| 6/17/2024 | SF2 | NA | NA |
| 6/17/2024 | R2 | 150 | NA |
| 6/25/2024 | SF2 | 83 | NA |
| 6/25/2024 | SF2 | 125 | 7.75 |
| 6/25/2024 | SF2 | 27 | 8.55 |
| 6/25/2024 | N1 | 120 | NA |
| 6/25/2024 | DP1 | 120 | 7.85 |
| 6/25/2024 | N4 | 87 | NA |
| 6/25/2024 | N6 | 61.2 | NA |
| 6/25/2024 | R2 | 148 | 7 |
| 7/9/2024 | SF2 | 133 | 7.59 |
| 7/9/2024 | SF2 | 137 | 7.47 |
| 7/9/2024 | SF2 | 28.9 | 8.4 |
| 7/9/2024 | R2 | 143.9 | 6.94 |
| 7/9/2024 | S5 | 127 | 6.56 |
| 7/9/2024 | R5 | 154 | 7.11 |
| 7/9/2024 | R4 | 54 | 8.56 |
| 7/16/2024 | A12 | 134.6 | 7.18 |
| 7/16/2024 | A11 | 23 | 8.78 |
| 7/16/2024 | A13 | 148.3 | 7.2 |

| | | | |
|---|---|---|---|
| 7/16/2024 | A14 | 20 | 8.3 |
| 7/16/2024 | A8 | 0.985 | 8.61 |
| 7/16/2024 | A4 | 29.1 | 8.47 |
| 7/25/2024 | E12 | 35.8 | 8.56 |
| 7/25/2024 | E12 | 36.9 | 8.72 |
| 7/25/2024 | E12 | 56 | 8.6 |
| 7/25/2024 | E12 | 63 | NA |
| 7/25/2024 | E13 | 63 | 8.25 |
| 7/25/2024 | E13 | 33.5 | 8.4 |
| 7/25/2024 | E13 | 43.9 | 9.1 |
| 7/25/2024 | E14 | 32 | 8.3 |
| 7/30/2024 | E14B | 54 | 8.6 |
| 7/30/2024 | E15B | 39.4 | 8.64 |
| 7/30/2024 | R2 | 140.4 | 7.03 |
| 8/12/2024 | A2W | 27.6 | 7.7 |
| 8/12/2024 | A2E | 33.2 | 9.56 |
| 8/12/2024 | AB1 | 29.4 | 8.63 |
| 8/12/2024 | SF2 | 137.1 | 7.18 |
| 8/12/2024 | SF2 | 59 | NA |
| 8/12/2024 | R2 | 138 | 7.14 |
| 8/20/2024 | N1 | 104.4 | 7.82 |
| 8/20/2024 | DP1 | 110 | 7.66 |
| 8/20/2024 | N2 | NA | NA |
| 8/20/2024 | N2 | 88.1 | 8.08 |
| 8/20/2024 | N3 | 81.2 | 8.2 |
| 8/20/2024 | N6 | 40.6 | 9.07 |
| 8/20/2024 | N4 | 63.5 | 8.68 |
| 8/20/2024 | R2 | 116.1 | 7.03 |
| 8/26/2024 | SF2 | 140.5 | 7.08 |
| 8/26/2024 | SF2 | 34.8 | 8.87 |
| 8/26/2024 | R2 | 127.2 | 6.95 |

**Online Resource 4.** Dated observations (including salinity and pH measurements) in identified SBSPRP ponds in which at least five living *Ephydra* (of any life stage) of a given species were found. "NA" indicates the recorded measurement was not taken, lost or illegible. LBS = Living Brine Shrimp, Sal = salinity (in ppt).

| Coll_Date | PondName | Notes | LBS | Sal | pH | Species |
|---|---|---|---|---|---|---|
| 5/23/2024 | SF2 (east) small pond | Note: SF2 (east) small pond corresponds to these approximate GPS coordinates: 37.497618, -122.129106 | TRUE | 104 | 7.66 | *gracilis* |
| 5/23/2024 | SF2 (east) bayside | Note: SF2 (east) bayside corresponds to these approximate GPS coordinates: 37.4973651223155, -122.12835945390223 | FALSE | 28 | 8.3 | *millbrae* |
| 5/23/2024 | R2 | Some flies. Full of brine shrimp. | TRUE | 96 | 7.14 | *gracilis* |
| 6/2/2024 | Kirby Park, Elkhorm Slough | Lots of flies, more in the seeps (approx 52 ppm), while 'waterside' was about 27 ppt. Lots of millbrae in the 52 ppt area, while there was another fly, a bit smaller, dull brown in the 'waterside' area. | FALSE | 52 | NA | *millbrae* |
| 6/17/2024 | SF2 (east) small pond | Bayside 28 ppt, small pond 115 ppt. No longer any brine shrimp in small pond. Also not a lot of adult flies. Most pupae I observed had already "popped off". Bayside area had a lot of Lipocheata ranica. | FALSE | 115 | NA | *gracilis* |
| 6/17/2024 | SF2 (east) bayside | Bayside 28 ppt, small pond 115 ppt. No longer any brine shrimp in small pond. Also not a lot of adult flies. Most pupae I observed had already "popped off". Bayside area had a lot of Lipocheata ranica. | FALSE | 28 | NA | *millbrae* |
| 6/17/2024 | R2 | 150 ppt, pH 7.01. Extremely saturated salt. Some very red brine shrimp but not nearly as abundant as previously. | TRUE | 150 | 7.01 | *gracilis* |
| 6/25/2024 | SF2 (west) | Lots of larvae and pupae; a few brine shrimp. Note: SF2 (west) orresponds to these approximate GPS coordinates: 37.497207649074475, -122.12862499259546 | TRUE | 83 | NA | *gracilis* |
| 6/25/2024 | SF2 (east) small pond | Lots of larvae and brine shrimp; larvae seemed to be quite big, approx L3. | TRUE | 125 | 7.75 | *gracilis* |
| 6/25/2024 | SF2 (east) bayside | Some adult flies. Did not observe any larvae or brine shrimp. Air bubbles forming on top of mats. | FALSE | 27 | 8.55 | *millbrae* |
| 6/25/2024 | N1 | Lots of flies along the shore. At southernmost end, some brine shrimp, and some larvae. More of these were found when the soil in the water was dark, as opposed to brown. | TRUE | 120 | NA | *gracilis* |
| 6/25/2024 | DP1 | Abundance of flies. | NA | 120 | 7.85 | *gracilis* |
| 6/25/2024 | R2 | Small, bright red brine shrimp. Some larvae (seem to be about L2). On the Ravenswood trail after Structural Collapse, was still 150 ppt, small amount of shrimp, some dead larvae. | TRUE | 148 | 7 | *gracilis* |
| 7/9/2024 | SF2 (west) | | FALSE | 133 | 7.59 | *gracilis* |
| 7/9/2024 | SF2 (east) small pond | Some very red, dead brine shrimp. Lots of L3 and pupae. | FALSE | 137 | 7.47 | *gracilis* |
| 7/9/2024 | SF2 (east) bayside | Determined how millbrae L and P look. Foubd in little ponds with lots of other life including water boatmen. | FALSE | 28.9 | 8.4 | *millbrae* |
| 7/9/2024 | R2 | Some bright red larvae, some L3 brine fly larvae, very small. Remarkably dried up compared to previously. Salt saturated. | TRUE | 143.9 | 6.94 | *gracilis* |

| | | | | | | |
|---|---|---|---|---|---|---|
| 7/9/2024 | R5 | Lots of small dark pupae and some L3. Some brine shrimp, which were not noticeably red. | TRUE | 154 | 7.11 | *gracilis* |
| 7/9/2024 | R4 | No brine shrimp. Flies caught in a puddle of 54 ppt, pH 8.56, with pickleweed and algae. In the larger R4 body of water, 31 ppt, 27C, pH 8.13. | FALSE | 54 | 8.56 | *millbrae* |
| 7/16/2024 | A11 | Some algae, greenish water. (Note: double checked these were indeed gracilis). | FALSE | 23 | 8.78 | *gracilis* |
| 7/16/2024 | A13 | Lots of brine shrimp, L3 and P; lots of flies. Some brine shrimp, bright-red. | TRUE | 148.3 | 7.2 | *gracilis* |
| 7/16/2024 | A14 | Some algae, greenish water.. (Note: double checked these were indeed gracilis). | FALSE | 20 | 8.3 | *gracilis* |
| 7/25/2024 | E12 a | Low salinity. Pickleweed. Note: E12a corresponds to approximate GPS coordinates: 37.614391384631084, -122.11848854105618 | FALSE | 35.8 | 8.56 | *millbrae* |
| 7/25/2024 | E12b | Lots of flies, but a few pupae or larvae. Lots of flies on microbial mats. Note: E12b corresponds to approximate GPS coordinates: 37.61575116436146, -122.12196468394936 | FALSE | 36.9 | 8.72 | *millbrae* |
| 7/25/2024 | E12c | Lots of little fish. Note: E12c corresponds to approximate GPS coordinates 37.61486731036518, -122.12436794323352 | FALSE | 56 | 8.6 | *millbrae* |
| 7/30/2024 | Hayward Shoreline | Small golden flies observed mating in puddle. (Note: genotyping later determined these were E. millbrae, albeit with striking coloration). | FALSE | 32.2 | NA | *millbrae* |
| 7/30/2024 | E14B | No brine shrimp. | FALSE | 54 | 8.6 | *millbrae* |
| 7/30/2024 | E15B | Pupae but no larvae or brine shrimp. many flies. | FALSE | 39.4 | 8.64 | *millbrae* |
| 7/30/2024 | R2 | Some larvae and brine shrmp, very small. | TRUE | 140.4 | 7.03 | *gracilis* |
| 8/12/2024 | SF2 (east) small pond | Lots of L3, a few pupae, some flies. No brine shrimp. | FALSE | 137.1 | 7.18 | *gracilis* |
| 8/12/2024 | SF2 (east) bayside | Lots of millbrae. | FALSE | 59 | NA | *millbrae* |
| 8/12/2024 | R2 | Some L3, some small red brine shrimp. | TRUE | 138 | 7.14 | *gracilis* |
| 8/20/2024 | N1 | Some diving beetles. P3 pupae sparse. Very few living brine shrimp. | TRUE | 104.4 | 7.82 | *gracilis* |
| 8/20/2024 | N2 | Water lime green. P3 pupae. | NA | 88.1 | 8.08 | *gracilis* |
| 08/26/2024 | Bair Island | Pond off of boardwalk. Very muddy. Water boat men, millbrae larvae. | FALSE | 44.5 | 8.66 | *millbrae* |
| 08/26/2024 | SF2 (small pond) | No visible brine shrimp. | FALSE | 140.5 | 7.08 | *gracilis* |
| 08/26/2024 | SF2 (bayside) | | FALSE | 34.8 | 8.87 | *millbrae* |
| 08/26/2024 | R2 | Some floating pupae. Reddish. | FALSE | 127.2 | 6.95 | *gracilis* |

**Online Resource 5.** % *COI* similarity for tips on **Fig 4** *Ephydra* gene genealogy.

| | PV790562 | PV790570 | KU496737..1 | JF874959.1 | EU493562.1 | MG081213.1 | JF867574.1 | MF059325.1 | MZ623564.1 | PV790571.1 | PQ772809 | PV790564.1 | PV790567 | PQ772810.1 | PQ772806.1 | OP971562.1 | OP971557.1 |
|---|---|---|---|---|---|---|---|---|---|---|---|---|---|---|---|---|---|
| PV790562.1 Unknown sp. Mono Lake, CA | | 99.73 | 99.83 | 99.73 | 99.75 | 99.64 | 99.67 | 98.65 | 98.73 | 98.4 | 98.89 | 94.57 | 93.69 | 93.26 | 93.49 | 93.91 | 94.51 |
| PV790570.1 Unknown sp. Great Salt Lake, UT | 99.73 | | 100 | 100 | 100 | 99.82 | 99.83 | 98.78 | 98.98 | 98.19 | 98.21 | 94.29 | 93.66 | 92.75 | 92.95 | 93.91 | 94.64 |
| KU496737.1 E. riparia Alaska | 99.83 | 100 | | 100 | 100 | 99.82 | 99.83 | 99.34 | 98.98 | 98.69 | 99.43 | 94.29 | 94.61 | 94.08 | 94.34 | 93.91 | 94.81 |
| JF874959.1 Ephydra sp. Manitoba, Canada | 99.73 | 100 | 100 | | 100 | 99.82 | 99.83 | 98.78 | 98.98 | 98.81 | 99.13 | 94.29 | 93.66 | 93.25 | 93.47 | 93.91 | 94.64 |
| EU493562.1 E. packardi | 99.75 | 100 | 100 | 100 | | 100 | 99.75 | 98.98 | 98.98 | 99.18 | 99.43 | 94.29 | 94.61 | 94.35 | 94.34 | 93.91 | 94.42 |
| MG081213.1 Ephydra sp. Ontario, Canada | 99.64 | 99.82 | 99.82 | 99.82 | 100 | | 99.64 | 99.09 | 99.18 | 98.59 | 99.38 | 94.21 | 94.44 | 93.88 | 94.17 | 93.97 | 94.72 |
| JF867574.1 E. riparia BOLD | 99.67 | 99.83 | 99.83 | 99.83 | 99.75 | 99.64 | | 99.17 | 98.73 | 98.44 | 99.17 | 94.02 | 94.34 | 93.83 | 94.09 | 93.65 | 94.64 |
| MF059325.1 E. macellaria Malta | 98.65 | 98.78 | 99.34 | 98.78 | 98.98 | 99.09 | 99.17 | | 98.48 | 97 | 97.28 | 94.57 | 94.63 | 94.16 | 94.41 | 94.42 | 94.64 |
| MZ623564.1 E. macellaria Finland | 98.73 | 98.98 | 98.98 | 98.98 | 98.98 | 99.18 | 98.73 | 98.48 | | 98.16 | 98.41 | 94.02 | 94.07 | 93.85 | 93.83 | 93.4 | 93.91 |
| PV790571.1 Riparia-like Ephydra sp. Great Salt Lake, UT | 98.4 | 98.19 | 98.69 | 98.81 | 99.18 | 98.59 | 98.44 | 97 | 98.16 | | 98.88 | 94.16 | 93.66 | 93.3 | 93.4 | 93.08 | 93.27 |
| PQ772809.1 E. riparia Russia | 98.89 | 98.21 | 99.43 | 99.13 | 99.43 | 99.38 | 99.17 | 97.28 | 98.41 | 98.88 | | 94.23 | 93.54 | 93 | 92.73 | 93.3 | 93.25 |
| PV790564.1 E. millbrae South Bay Salt Ponds, CA | 94.57 | 94.29 | 94.29 | 94.29 | 94.29 | 94.21 | 94.02 | 94.57 | 94.02 | 94.16 | 94.23 | | 99.71 | 99.52 | 99.73 | 94.02 | 94.57 |
| PV790567 E. millbrae South Bay Salt Ponds, CA | 93.69 | 93.66 | 94.61 | 93.66 | 94.61 | 94.44 | 94.34 | 94.63 | 94.07 | 93.66 | 93.54 | 99.71 | | 100 | 99.76 | 93.53 | 93.59 |
| PQ772810.1 E. millbrae Hayward, CA | 93.26 | 92.75 | 94.08 | 93.25 | 94.35 | 93.88 | 93.83 | 94.16 | 93.85 | 93.3 | 93 | 99.52 | 100 | | 99.11 | 93.34 | 93.27 |
| PQ772806.1 E. millbrae South Bay Salt Ponds, CA | 93.49 | 92.95 | 94.34 | 93.47 | 94.34 | 94.17 | 94.09 | 94.41 | 93.83 | 93.4 | 92.73 | 99.73 | 99.76 | 99.11 | | 93.57 | 93.63 |
| OP971562.1 E. gracilis Great Salt Lake, UT | 93.91 | 93.91 | 93.91 | 93.91 | 93.91 | 93.97 | 93.65 | 94.42 | 93.4 | 93.08 | 93.3 | 94.02 | 93.53 | 93.34 | 93.57 | | 99.24 |
| OP971557.1 E. gracilis Great Salt Lake, UT | 94.51 | 94.64 | 94.81 | 94.64 | 94.42 | 94.72 | 94.64 | 94.64 | 93.91 | 93.27 | 93.25 | 94.57 | 93.59 | 93.27 | 93.63 | 99.24 | |
| JF867420.1 Ephydridae sp. | 90.93 | 91.19 | 91.06 | 91.19 | 91.12 | 90.58 | 90.89 | 91.34 | 91.62 | 90.31 | 90.34 | 91.03 | 91.22 | 90.76 | 91.38 | 93.65 | 93.3 |
| PQ772805.1 E. gracilis South Bay Salt Ponds, CA | 87.15 | 87.08 | 87.15 | 87.41 | 87.15 | 86.94 | 86.89 | 87.65 | 87.66 | 87.64 | 86.69 | 88.22 | 87.8 | 88.33 | 88.28 | 92.54 | 93.14 |
| PQ772812.1 E. gracilis South Bay Salt Ponds, CA | 83.7 | 83.89 | 83.36 | 84.01 | 83.36 | 82.82 | 83.08 | 84.25 | 83.9 | 84.36 | 83.57 | 83.94 | 84.46 | 84.85 | 84.85 | 89.06 | 89.86 |
| PV790568.1 E. gracilis South Bay Salt Ponds, CA | 83.37 | 83.37 | 84.1 | 83.37 | 84.52 | 85.82 | 83.85 | 83.87 | 84.77 | 83.87 | 84.19 | 88.72 | 85.44 | 85.22 | 85.6 | 89.85 | 89.04 |
| PQ772817.1 E. gracilis South Bay Salt Ponds, CA | 85.49 | 85.49 | 85.49 | 85.49 | 85.49 | 85.07 | 85.22 | 86.02 | 86.02 | 85.62 | 85.62 | 86.94 | 88.52 | 87.34 | 87.2 | 90.24 | 89.97 |
| PV790563.1 E. hians Lake Abert, OR | 87.91 | 87.66 | 88.01 | 87.66 | 88.01 | 88.15 | 87.76 | 87.66 | 88.52 | 87.53 | 87.59 | 88.32 | 88.03 | 87.97 | 88.2 | 90.82 | 90.68 |
| OP971547.1 E. hians Great Salt Lake, UT | 89.32 | 89.28 | 89.79 | 89.28 | 88.58 | 89.98 | 89.62 | 89.28 | 88.58 | 87.68 | 87.41 | 88.59 | 87.69 | 87.44 | 87.87 | 91.37 | 91.46 |
| HM374264.1 E. hians BOLD | 89.24 | 89.21 | 89.57 | 89.21 | 88.32 | 89.31 | 89.4 | 89.36 | 88.32 | 87.59 | 87.33 | 88.32 | 87.8 | 87.59 | 88 | 91.12 | 91.29 |
| PQ772816.1 E. hians Lee Vining, CA | 88.47 | 88.22 | 88.58 | 88.22 | 88.58 | 88.49 | 88.32 | 88.22 | 88.32 | 87.91 | 88.1 | 88.59 | 88.03 | 87.91 | 88.2 | 91.37 | 91.23 |
| PV790569 E. hians Great Salt Lake, UT | 87.38 | 87.13 | 88.17 | 87.41 | 88.17 | 88.33 | 87.92 | 88.11 | 88.17 | 88.04 | 87.13 | 88.77 | 88.54 | 88.71 | 88.88 | 91 | 90.44 |
| PV790572 E. hians Lake Abert, OR | 87.68 | 87.62 | 88.51 | 87.62 | 88.51 | 88.7 | 88.25 | 88.33 | 88.51 | 87.62 | 87.26 | 89.14 | 88.48 | 88.81 | 88.81 | 91.38 | 90.8 |
| PV790573.1 E. hians Mono Lake, CA | 89.24 | 89.56 | 89.74 | 89.21 | 88.32 | 89.67 | 89.57 | 89.67 | 88.32 | 87.09 | 86.55 | 88.59 | 88.54 | 87.91 | 88.32 | 91.12 | 90.79 |
| PQ772813.1 E. hians Great Salt Lake, UT | 88.09 | 87.81 | 88.43 | 88.11 | 88.43 | 88.33 | 88.17 | 87.88 | 88.17 | 88.71 | 87.81 | 88.77 | 87.8 | 88.04 | 88.21 | 91.26 | 91.18 |
| PQ772814.1 E. hians Great Salt Lake, UT | 87.62 | 87.36 | 88.17 | 87.65 | 88.17 | 88.06 | 87.92 | 87.88 | 88.69 | 88.26 | 87.36 | 88.49 | 87.8 | 88.04 | 88.21 | 91 | 90.69 |
| PQ772808.1 E. hians Deep Springs, CA | 86.04 | 86.04 | 86.21 | 86.04 | 86.21 | 86.21 | 85.91 | 85.78 | 86.21 | 86.04 | 85.65 | 87.06 | 86.57 | 86.57 | 86.57 | 88.86 | 88.9 |
| PQ772815.1 E. hians Owens Lake, CA | 87.31 | 87.06 | 87.06 | 87.06 | 87.06 | 87.12 | 86.8 | 86.8 | 87.56 | 86.74 | 86.93 | 87.23 | 87.74 | 87.25 | 87.53 | 89.85 | 90.1 |
| PQ772807.1 E. hians Big Soda Lake, CO | 84.06 | 83.81 | 84.31 | 83.81 | 84.31 | 84.63 | 84.05 | 83.56 | 84.05 | 83.5 | 83.67 | 84.15 | 83.62 | 83.5 | 83.61 | 86.84 | 86.57 |
| PV790574.1 Unknown sp., Walker Lake, Nevada | 90.45 | 90.28 | 90.44 | 90.28 | 89.85 | 90.04 | 90.27 | 90.62 | 89.85 | 88.74 | 88.99 | 89.4 | 89.52 | 89.49 | 89.49 | 91.12 | 92.06 |
| PZ634866.1 Riparia-like sp GSL UT | 88.92 | 88.02 | 88.95 | 88.02 | 90.47 | 89.58 | 88.79 | 88.02 | 90.47 | 89.29 | 89.83 | 89.27 | 89.31 | 89.04 | 89.02 | 88.9 | 88.1 |
| AB669723.1 Drosophila punjabensis | 87.08 | 87.35 | 87.25 | 87.08 | 85.28 | 86.41 | 87.09 | 87.39 | 84.77 | 84.56 | 83.86 | 85.05 | 85.61 | 85.49 | 85.5 | 88.32 | 89.28 |
| KU875234.1 Hydrellia griseola Dipt outgroup | 86.15 | 86.02 | 86.42 | 86.02 | 85.79 | 86.23 | 86.26 | 86.32 | 85.79 | 84.86 | 84.78 | 86.96 | 87.32 | 87.02 | 87.3 | 86.55 | 87.77 |
| GU673025.1 Maniola jurtina - moth | 86.3 | 86.17 | 86.42 | 86.17 | 87.56 | 85.51 | 86.26 | 87.39 | 88.07 | 86.9 | 86.63 | 86.14 | 86.59 | 86.68 | 86.95 | 89.59 | 88.11 |

| | JF867420.1 | PQ772805.1 | PQ772812.1 | PV790568.1 | PQ772817.1 | PV790563.1 | OP971547.1 | HM374264.1 | PQ772816.1 | PV790569 | PV790572 | PV790573.1 | PQ772813.1 | PQ772814.1 | PQ772808.1 | PQ772815.1 | PQ772807.1 | PV790574.1 | PZ634866.1 | AB669723.1 | KU875234.1 | GU673025.1 |
|---|---|---|---|---|---|---|---|---|---|---|---|---|---|---|---|---|---|---|---|---|---|---|
| PV790562.1 Unknown sp. Mono Lake, CA | 90.93 | 87.15 | 83.7 | 83.37 | 85.49 | 87.91 | 89.32 | 89.24 | 88.47 | 87.38 | 87.68 | 89.24 | 88.09 | 87.62 | 86.04 | 87.31 | 84.06 | 90.45 | 88.92 | 87.08 | 86.15 | 86.3 |
| PV790570.1 Unknown sp. Great Salt Lake, UT | 91.19 | 87.08 | 83.89 | 83.37 | 85.49 | 87.66 | 89.28 | 89.21 | 88.22 | 87.13 | 87.62 | 89.56 | 87.81 | 87.36 | 86.04 | 87.06 | 83.81 | 90.28 | 88.02 | 87.35 | 86.02 | 86.17 |
| KU496737.1 E. riparia Alaska | 91.06 | 87.15 | 83.36 | 84.1 | 85.49 | 88.01 | 89.79 | 89.57 | 88.58 | 88.17 | 88.51 | 89.74 | 88.43 | 88.17 | 86.21 | 87.06 | 84.31 | 90.44 | 88.95 | 87.25 | 86.42 | 86.42 |
| JF874959.1 Ephydra sp. Manitoba, Canada | 91.19 | 87.41 | 84.01 | 83.37 | 85.49 | 87.66 | 89.28 | 89.21 | 88.22 | 87.41 | 87.62 | 89.21 | 88.11 | 87.65 | 86.04 | 87.06 | 83.81 | 90.28 | 88.02 | 87.08 | 86.02 | 86.17 |
| EU493562.1 E. packardi | 91.12 | 87.15 | 83.36 | 84.52 | 85.49 | 88.01 | 88.58 | 88.32 | 88.58 | 88.17 | 88.51 | 88.32 | 88.43 | 88.17 | 86.21 | 87.06 | 84.31 | 89.85 | 90.47 | 85.28 | 85.79 | 87.56 |
| MG081213.1 Ephydra sp. Ontario, Canada | 90.58 | 86.94 | 82.82 | 85.82 | 85.07 | 88.15 | 89.98 | 89.31 | 88.49 | 88.33 | 88.7 | 89.67 | 88.33 | 88.06 | 86.21 | 87.12 | 84.63 | 90.04 | 89.58 | 86.41 | 86.23 | 85.51 |
| JF867574.1 E. riparia BOLD | 90.89 | 86.89 | 83.08 | 83.85 | 85.22 | 87.76 | 89.62 | 89.4 | 88.32 | 87.92 | 88.25 | 89.57 | 88.17 | 87.92 | 85.91 | 86.8 | 84.05 | 90.27 | 88.79 | 87.09 | 86.26 | 86.26 |
| MF059325.1 E. macellaria Malta | 91.34 | 87.65 | 84.25 | 83.87 | 86.02 | 87.66 | 89.28 | 89.36 | 88.22 | 88.11 | 88.33 | 89.67 | 87.88 | 87.88 | 85.78 | 86.8 | 83.56 | 90.62 | 88.02 | 87.39 | 86.32 | 87.39 |
| MZ623564.1 E. macellaria Finland | 91.62 | 87.66 | 83.9 | 84.77 | 86.02 | 88.52 | 88.58 | 88.32 | 88.32 | 88.17 | 88.51 | 88.32 | 88.17 | 88.69 | 86.21 | 87.56 | 84.05 | 89.85 | 90.47 | 84.77 | 85.79 | 88.07 |
| PV790571.1 Riparia-like Ephydra sp. Great Salt Lake, UT | 90.31 | 87.64 | 84.36 | 83.87 | 85.62 | 87.53 | 87.68 | 87.59 | 87.91 | 88.04 | 87.62 | 87.09 | 88.71 | 88.26 | 86.04 | 86.74 | 83.5 | 88.74 | 89.29 | 84.56 | 84.86 | 86.9 |
| PQ772809.1 E. riparia Russia | 90.34 | 86.69 | 83.57 | 84.19 | 85.62 | 87.59 | 87.41 | 87.33 | 88.1 | 87.13 | 87.26 | 86.55 | 87.81 | 87.36 | 85.65 | 86.93 | 83.67 | 88.99 | 89.83 | 83.86 | 84.78 | 86.63 |
| PV790564.1 E. millbrae South Bay Salt Ponds, CA | 91.03 | 88.22 | 83.94 | 88.72 | 86.94 | 88.32 | 88.59 | 88.32 | 88.59 | 88.77 | 89.14 | 88.59 | 88.77 | 88.49 | 87.06 | 87.23 | 84.15 | 89.4 | 89.27 | 85.05 | 86.96 | 86.14 |
| PV790567 E. millbrae South Bay Salt Ponds, CA | 91.22 | 87.8 | 84.46 | 85.44 | 88.52 | 88.03 | 87.69 | 87.8 | 88.03 | 88.54 | 88.48 | 88.54 | 87.8 | 87.8 | 86.57 | 87.74 | 83.62 | 89.52 | 89.31 | 85.61 | 87.32 | 86.59 |
| PQ772810.1 E. millbrae Hayward, CA | 90.76 | 88.33 | 84.85 | 85.22 | 87.34 | 87.97 | 87.44 | 87.59 | 87.91 | 88.71 | 88.81 | 87.91 | 88.04 | 88.04 | 86.57 | 87.25 | 83.5 | 89.49 | 89.04 | 85.49 | 87.02 | 86.68 |
| PQ772806.1 E. millbrae South Bay Salt Ponds, CA | 91.38 | 88.28 | 84.85 | 85.6 | 87.2 | 88.2 | 87.87 | 88 | 88.2 | 88.88 | 88.81 | 88.32 | 88.21 | 88.21 | 86.57 | 87.53 | 83.61 | 89.49 | 89.02 | 85.5 | 87.3 | 86.95 |
| OP971562.1 E. gracilis Great Salt Lake, UT | 93.65 | 92.54 | 89.06 | 89.85 | 90.24 | 90.82 | 91.37 | 91.12 | 91.37 | 91 | 91.38 | 91.12 | 91.26 | 91 | 88.86 | 89.85 | 86.84 | 91.12 | 88.9 | 88.32 | 86.55 | 89.59 |
| OP971557.1 E. gracilis Great Salt Lake, UT | 93.3 | 93.14 | 89.86 | 89.04 | 89.97 | 90.68 | 91.46 | 91.29 | 91.23 | 90.44 | 90.8 | 90.79 | 91.18 | 90.69 | 88.9 | 90.1 | 86.57 | 92.06 | 88.1 | 89.28 | 87.77 | 88.11 |
| JF867420.1 Ephydridae sp. | | 90.91 | 87.93 | 86.33 | 88.13 | 90.43 | 90.79 | 91.03 | 90.23 | 90.68 | 90.71 | 91.03 | 90.68 | 90.44 | 88.69 | 89.85 | 85.07 | 89.95 | 85.37 | 87.99 | 87.84 | 87.84 |
| PQ772805.1 E. gracilis South Bay Salt Ponds, CA | 90.91 | | 96.5 | 95.62 | 94.46 | 85.53 | 86.03 | 86.48 | 85.79 | 86.63 | 86.43 | 85.95 | 87.08 | 86.85 | 85.52 | 84.58 | 81.45 | 87.95 | 84.79 | 87.75 | 86.71 | 88.34 |
| PQ772812.1 E. gracilis South Bay Salt Ponds, CA | 87.93 | 96.5 | | 92.23 | 92.49 | 81.97 | 82.62 | 83.27 | 82.24 | 83.41 | 82.9 | 82.94 | 83.89 | 83.65 | 85.52 | 81.59 | 78.33 | 84.21 | 80.74 | 84.83 | 83.27 | 85.48 |
| PV790568.1 E. gracilis South Bay Salt Ponds, CA | 86.33 | 95.62 | 92.23 | | 93.14 | 83.31 | 83.37 | 83.13 | 83.02 | 83.69 | 83.89 | 83.37 | 83.44 | 83.44 | 82.7 | 82.49 | 79.05 | 85.7 | 83.59 | 84.61 | 83.37 | 83.62 |
| PQ772817.1 E. gracilis South Bay Salt Ponds, CA | 88.13 | 94.46 | 92.49 | 93.14 | | 85.22 | 85.49 | 85.22 | 85.22 | 85.49 | 85.45 | 85.49 | 85.22 | 85.22 | 85.4 | 84.3 | 81.57 | 86.28 | 84.05 | 86.81 | 85.22 | 85.75 |
| PV790563.1 E. hians Lake Abert, OR | 90.43 | 85.53 | 81.97 | 83.31 | 85.22 | | 99.5 | 99.24 | 98.24 | 97.72 | 98.45 | 98.49 | 97.97 | 99.49 | 98.33 | 98.09 | 91.54 | 87.79 | 84.38 | 85.89 | 85.14 | 85.64 |
| OP971547.1 E. hians Great Salt Lake, UT | 90.79 | 86.03 | 82.62 | 83.37 | 85.49 | 99.5 | | 99.5 | 98.75 | 97.55 | 98.26 | 98.16 | 98.28 | 99.51 | 98.39 | 97.59 | 92.08 | 88.77 | 84.92 | 87.44 | 86.77 | 86.1 |
| HM374264.1 E. hians BOLD | 91.03 | 86.48 | 83.27 | 83.13 | 85.22 | 99.24 | 99.5 | | 99 | 97.9 | 98.57 | 98.33 | 98.6 | 99.3 | 98.48 | 97.34 | 92.34 | 89.28 | 83.22 | 87.84 | 86.17 | 86.47 |
| PQ772816.1 E. hians Lee Vining, CA | 90.23 | 85.79 | 82.24 | 83.02 | 85.22 | 98.24 | 98.75 | 99 | | 98.48 | 99.23 | 99.25 | 99.75 | 98.22 | 97.46 | 96.95 | 92.84 | 87.34 | 84.46 | 85.21 | 84.96 | 86.22 |
| PV790569 E. hians Great Salt Lake, UT | 90.68 | 86.63 | 83.41 | 83.69 | 85.49 | 97.72 | 97.55 | 97.9 | 98.48 | | 99.52 | 98.65 | 98.31 | 97.63 | 97.42 | 96.4 | 91.48 | 87.69 | 84.52 | 85.89 | 85.55 | 87.41 |
| PV790572 E. hians Lake Abert, OR | 90.71 | 86.43 | 82.9 | 83.89 | 85.45 | 98.45 | 98.26 | 98.57 | 99.23 | 99.52 | | 100 | 98.81 | 98.1 | 97.41 | 97.13 | 92.25 | 88.02 | 85.08 | 86.19 | 85.48 | 87.38 |
| PV790573.1 E. hians Mono Lake, CA | 91.03 | 85.95 | 82.94 | 83.37 | 85.49 | 98.49 | 98.16 | 98.33 | 99.25 | 98.65 | 100 | | 97.97 | 97.29 | 97.42 | 97.21 | 92.34 | 89.28 | 83.22 | 88.24 | 86.32 | 86.93 |
| PQ772813.1 E. hians Great Salt Lake, UT | 90.68 | 87.08 | 83.89 | 83.44 | 85.22 | 97.97 | 98.28 | 98.6 | 99.75 | 98.31 | 98.81 | 97.97 | | 98.31 | 97.16 | 96.66 | 92.49 | 87.18 | 84.26 | 85.21 | 84.85 | 86.71 |
| PQ772814.1 E. hians Great Salt Lake, UT | 90.44 | 86.85 | 83.65 | 83.44 | 85.22 | 99.49 | 99.51 | 99.3 | 98.22 | 97.63 | 98.1 | 97.29 | 98.31 | | 97.95 | 97.56 | 91.48 | 87.44 | 84.26 | 85.67 | 85.08 | 86.71 |
| PQ772808.1 E. hians Deep Springs, CA | 88.69 | 85.52 | 85.52 | 82.7 | 85.4 | 98.33 | 98.39 | 98.48 | 97.46 | 97.42 | 97.41 | 97.42 | 97.16 | 97.95 | | 96.98 | 90.6 | 87.13 | 82.24 | 84.72 | 83.4 | 84.72 |
| PQ772815.1 E. hians Owens Lake, CA | 89.85 | 84.58 | 81.59 | 82.49 | 84.3 | 98.09 | 97.59 | 97.34 | 96.95 | 96.4 | 97.13 | 97.21 | 96.66 | 97.56 | 96.98 | | 91.16 | 86.68 | 83.42 | 84.64 | 84.26 | 84.77 |
| PQ772807.1 E. hians Big Soda Lake, CO | 85.07 | 81.45 | 78.33 | 79.05 | 81.57 | 91.54 | 92.08 | 92.34 | 92.84 | 91.48 | 92.25 | 92.34 | 92.49 | 91.48 | 90.6 | 91.16 | | 83.02 | 80.72 | 80.72 | 80.93 | 81.81 |
| PV790574.1 Unknown sp., Walker Lake, Nevada | 89.95 | 87.95 | 84.21 | 85.7 | 86.28 | 87.79 | 88.77 | 89.28 | 87.34 | 87.69 | 88.02 | 89.28 | 87.18 | 87.44 | 87.13 | 86.68 | 83.02 | | 84.02 | 88.78 | 87.44 | 87.77 |
| PZ634866.1 Riparia-like sp GSL UT | 85.37 | 84.79 | 80.74 | 83.59 | 84.05 | 84.38 | 84.92 | 83.22 | 84.46 | 84.52 | 85.08 | 83.22 | 84.26 | 84.26 | 82.24 | 83.42 | 80.72 | 84.02 | | 81.4 | 80.91 | 81.9 |
| AB669723.1 Drosophila punjabensis | 87.99 | 87.75 | 84.83 | 84.61 | 86.81 | 85.89 | 87.44 | 87.84 | 85.21 | 85.89 | 86.19 | 88.24 | 85.21 | 85.67 | 84.72 | 84.64 | 80.72 | 88.78 | 81.4 | | 86.63 | 87.84 |
| KU875234.1 Hydrellia griseola Dipt outgroup | 87.84 | 86.71 | 83.27 | 83.37 | 85.22 | 85.14 | 86.77 | 86.17 | 84.96 | 85.55 | 85.48 | 86.32 | 84.85 | 85.08 | 83.4 | 84.26 | 80.93 | 87.44 | 80.91 | 86.63 | | 89.06 |
| GU673025.1 Maniola jurtina - moth | 87.84 | 88.34 | 85.48 | 83.62 | 85.75 | 85.64 | 86.1 | 86.47 | 86.22 | 87.41 | 87.38 | 86.93 | 86.71 | 86.71 | 84.72 | 84.77 | 81.81 | 87.77 | 81.9 | 87.84 | 89.06 | |
